# Phosphoproteomics of Hypertrophic Cardiomyopathy Patient Myocardium and Novel hiPSC-CM Model Reveal Protein Kinase A as a Modulator of Microtubule Repolymerization

**DOI:** 10.64898/2026.02.18.706710

**Authors:** Sıla Algül, Inez Duursma, Jennifer Hesson, Julie Mathieu, Richard de Goeij-de Haas, Alex A. Henneman, Sander R. Piersma, Thang V. Pham, Stephan A.C. Schoonvelde, Michelle Michels, Jean-Marc Soleilhac, Marie-Jo Moutin, Connie R. Jimenez, Michael Regnier, Diederik W.D. Kuster, Jolanda van der Velden

## Abstract

**Background and aims:** Increased levels of α-tubulin and its post-translational modifications (PTMs) are found in human heart failure and could initiate diastolic dysfunction by modulating cardiomyocyte stiffness. How these modifications occur and how they may underlie cardiac dysfunction remains unknown. Upstream kinases may play a critical role, but this has not been explored.

**Methods and results:** Here we address this question by, for the first time ever, determining levels of the enzymes involved in microtubule (MT) detyrosination and acetylation (αTAT1, HDAC6) in a well-characterized cohort of patients with hypertrophic cardiomyopathy (HCM). In HCM patients (N=10-11), protein levels of detyrosination enzymes remain unaltered, whilst levels of αTAT1 and HDAC6 were decreased and increased, respectively. Phosphoproteomics in HCM (N=24) and control (N=8) myocardium identified significant differences in over 1900 serine/threonine and 160 tyrosine phosphosites, in addition to increased EGFR/IGF1R-MAPK signaling in HCM. We subsequently showed that MT repolymerization was increased in HCM *MYBPC3*_Arg943X_ hiPSC-CMs. Isoprenaline-mediated PKA activation decreased MT repolymerization in hiPSC-CMs and revealed *CLASP1*, *MAST4* and *MAP1A* as potential MT modifiers in HCM.

**Conclusions:** We show that the altered HCM MT code cannot be attributed to levels of key MT-modifying enzymes. By combining kinome analyses in human HCM hearts with hiPSC-CM studies on MT dynamics, PTMs and contractility we unveiled a regulatory role for MTs in the cardiomyocyte response to beta-adrenergic receptor stimulation. Disease-mediated changes in the MT code thereby exert both a direct, and indirect effect on cardiac function via mediating the response to adrenergic activation.

**Graphical Abstract** created with BioRender.com

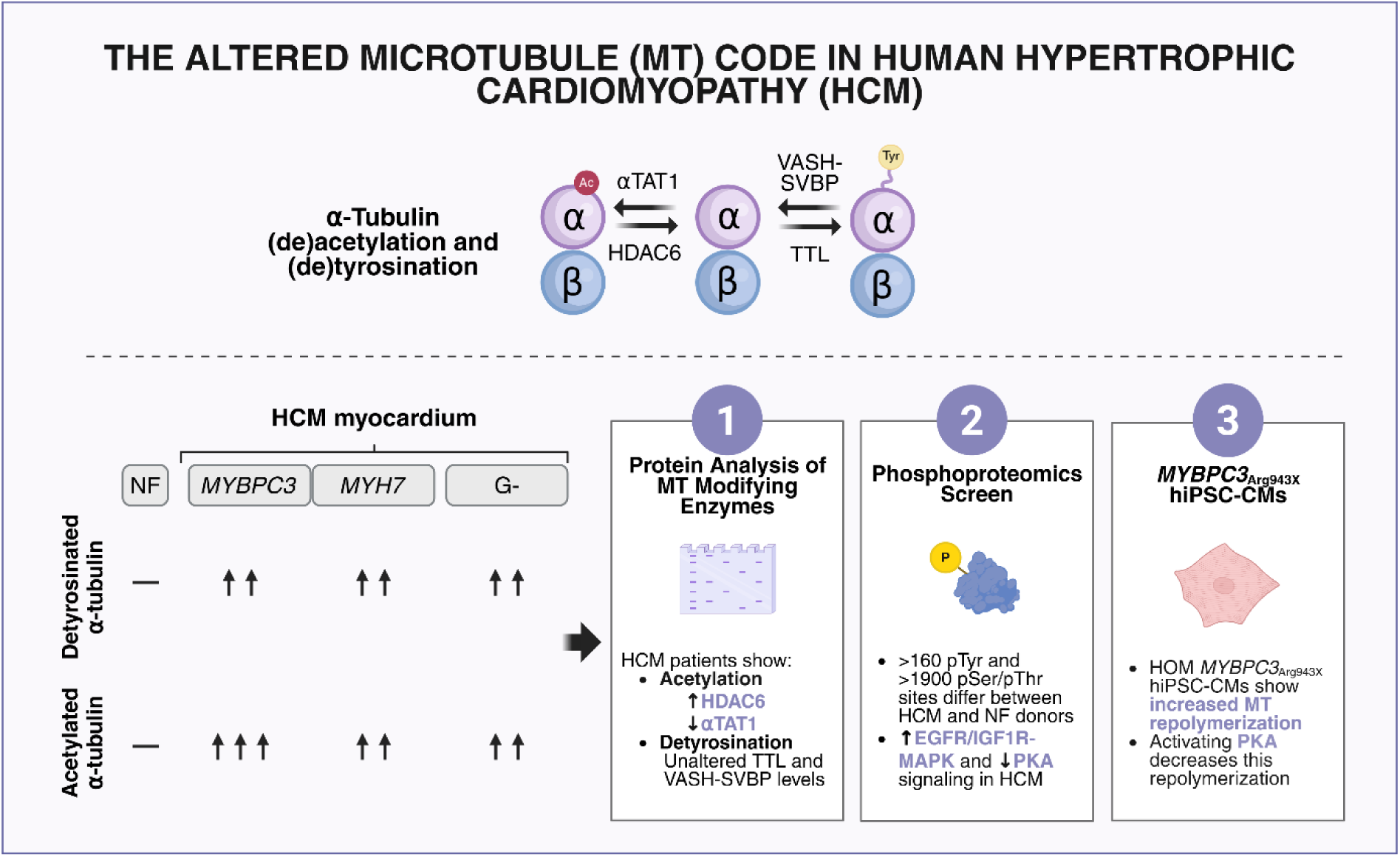

## Introduction

Hypertrophic cardiomyopathy (HCM) is characterized by hypertrophy of the muscular walls of the heart, which is accompanied by diastolic dysfunction and a preserved or even slightly elevated left ventricular (LV) ejection fraction. In approximately half of all patients, HCM is caused by a heterozygous pathogenic variant, i.e. mutation, in one of the genes encoding sarcomere proteins [1]. These patients are referred to as genotype-positive (G+) [2]. Impaired diastolic function can already present at the asymptomatic preclinical stage of the disease in mutation-carriers, which has been ascribed to mutation-mediated changes in sarcomere function and altered calcium handling [3, 4]. In the remaining half of the HCM patient population the actual cause of cardiac hypertrophy and impaired relaxation is unknown as no gene defect is identified, marking them as genotype-negative (G-) [1, 2].

In both G+ and G- HCM patients structural remodeling of the heart, including asymmetric LV hypertrophy and fibrosis, contributes to impaired relaxation during disease progression. At the level of the cardiomyocyte, a complex interplay of pathomechanisms underlie impaired muscle relaxation [5, 6]. Cardiomyocytes of heart failure patients with a preserved ejection fraction were shown to present with a titin-mediated increase in passive force development [7]. However, there is no evidence for such an increase in HCM [8, 9]. Instead, HCM patients present with impaired calcium handling, a decrease in the super-relaxed state of myosin, and dysregulated kinase signaling [10–13]. Recently, we have also found a significantly altered microtubule (MT) code in both G+ and G- HCM patients, which can contribute to relaxation deficits [14–17]. At the time of myectomy, HCM hearts consistently exhibit significantly elevated levels of α-tubulin and its post-translational modifications (PTMs), detyrosination and acetylation, compared to non-failing controls [14, 18]. Cleavage of the C-terminal tyrosine on α-tubulin results in detyrosination, a PTM that is mediated by vasohibin (VASH1 or VASH2) and a small vasohibin-binding protein (SVBP) complex. This PTM provides cardiomyocytes with resistance during contraction [19–21]. Though less studied, acetylation of α-tubulin on Lys40 by tubulin acetyltransferase 1 (αTAT1) is also posited to increase this viscoelastic resistance, which contributes to impaired muscle relaxation [22–24]. The ability of such PTMs to impair cardiomyocyte relaxation raises the question of how their increased abundance is mediated in HCM.

Intriguingly, when comparing genotypes, HCM myocardium with truncating *MYBPC3* mutations that cause haploinsufficiency of cardiac myosin-binding protein-C (cMyBP-C), demonstrates significantly higher levels of MT PTMs when compared to other genotypes (*MYH7* and thin filament mutations) and G- hearts [18]. Altered protein expression or regulation of the enzymes involved in detyrosination and acetylation could account for this difference. As key upstream enzymes modulate levels of MT detyrosination and acetylation, fine-tuning the levels of these enzymes has emerged as an important potential treatment strategy [15–17, 20, 25]. Yet, levels of these enzymes in failing myocardium have never been explored. In addition, previous human studies have been frequently performed in explanted end-stage failing hearts (NYHA class IV) with limited demographic and clinical data. [17]. It also remains to be determined whether more stable MTs precede and, perhaps, mediate diastolic dysfunction in HCM [26].

Thus, our aim was two-fold: (1) to unravel the upstream regulators of microtubule detyrosination and acetylation and (2) to dissect the involvement of MTs in the onset and early progression of cardiomyocyte dysfunction. To do so, we provide the first study of levels of key enzymes involved in MT detyrosination and acetylation in a unique and well-characterized HCM cohort with diastolic dysfunction and a preserved ejection fraction. We then combined this with a phosphoproteomics screen to assess the serine/threonine and tyrosine signature in myocardium of both G+ (*MYBPC3* and *MYH7*) and G- HCM patients for comparison to each other and to non-failing (NF) controls. In doing so, we screened for novel candidates that may alter MT dynamics in HCM and reveal possible causes of diastolic dysfunction in human cardiac disease. To address our second aim, we developed an HCM-associated induced-pluripotent stem cell-derived cardiomyocyte (hiPSC-CM) model of the Dutch founder variant *MYBPC3*_Arg943X_, carrying an α-tubulin bound red fluorescent protein (RFP) tag [27]. Ultimately, we employed this model to validate the involvement of kinases and phosphosites we found in our human HCM tissue screen.

## Methods

Expanded methods detailing protocols for human myocardial analysis, hiPSC-CM line generation and maintenance, contractility analyses, live cell imaging, and remaining assays can be found in the Supplemental Material.

### Human myocardium

All human tissue samples were obtained after written informed consent from each patient prior to surgery and from the patients’ or donors’ next of kin. For the human phosphoproteomics screen, we collected interventricular septal cardiac tissue from adult patients with obstructive HCM (N = 24; 15 males, 9 females, see **Table 1**) and NF controls (N = 8; 3 males, 5 females). All human studies complied with the Declaration of Helsinki.

**Table 1.** Cardiac characteristics of the HCM*_MYBPC3_*, HCM*_MYH7_* and G- patients.

|  | HCM<sub>MYBPC3</sub> | HCM<sub>MYH7</sub> | G- |
| --- | --- | --- | --- |
| Sex, male | 62.5% (5) | 62.5% (5) | 62.5% (5) |
| Age at myectomy, y | 45.0 ± 5.3 | 51.3 ± 5.1 | 60.0 ± 3.9 |
| MT code (values normalized to NF = 1). |  |  |  |
| α-tubulin/GAPDH | 7.4 ± 1.2 | 4.5 ± 0.6 | 3.6 ± 0.6* |
| Detyrosinated α-tubulin/GAPDH | 3.9 ± 0.5 | 3.9 ± 0.5 | 1.9 ± 0.3* |
| Acetylated α-tubulin/GAPDH | 12.3 ± 2.6 | 8.2 ± 2.1 | 4.6 ± 0.7* |
| Dimensions, mm |  |  |  |
| IVS | 21.6 ± 0.5 | 18.5 ± 0.9 | 18.1 ± 0.8** |
| IVSi | 10.3 ± 0.5 | 9.5 ± 1.2 | 8.4 ± 0.4* |
| LAD | 45.1 ± 1.4 | 49.7 ± 2.6 | 49.8 ± 1.3 |
| LADi | 22.2 ± 0.7 | 25.1 ± 0.9 | 24.9 ± 1.0 |
| EDD | 45.4 ± 2.2 | 43.0 ± 2.6 | 40.0 ± 3.2 |
| ESD | 22.5 ± 2.8 | na | na |
| Diastolic parameters |  |  |  |
| E/A ratio | 1.4 [0.8-1.9] | 1.5 [0.6-2.5] | 1.1 [0.5-1.7] |
| E/e' ratio | 18.6 [12.3-24.8] | 17.7 [10.9-24.6] | 20.7 [15.3-26.0] |
| TR velocity<br>(cm/s) | 2.3 [1.5-3.1] | 2.1 [1.6-2.7] | 2.8 [2.1-3.6] |
| Gradient at rest |  |  |  |
| LVOT<sub>g</sub> (mmHg) | 50.6 ± 11.7 | 61.6 ± 9.0 | 92.5 ± 16.6 |

\* $P < 0.05$ compared to HCM<sub>MYBPC3</sub>, \*\* $P < 0.01$ compared to HCM<sub>MYBPC3</sub>; na, not available.

### MYBPC3_Arg943X_ hiPSC-CM model

A parental hiPSC cell line carrying a mono-allelic mTag-RFP-T-TUBA1B (tubulin alpha 1b; clone AICS-0031-035; Allen Institute for Cell Science, Seattle, WA) was employed to generate wild-type (WT) and homozygous (HOM) *MYBPC3*_Arg943X_ clones using CRISPR/Cas9 technology, as described previously [33].

### Pharmacological interventions

WT and HOM hiPSC-CMs from at least 3 independently differentiated batches were treated with DMSO (0.1 vol.%, control, Sigma Aldrich, 472301), epoY (20 µM, detyrosination inhibitor, SML2301, Sigma-Aldrich), tubacin (HDAC6 inhibitor, Sigma Aldrich, SML0065, 10 µM), isoprenaline (100 nM, PKA agonist, Sigma Aldrich, I5627) or AEE 788 (0.01 µM, EGFR inhibitor, Tocris, 5318) for 1-2 hours (37°C).

### Data availability

Both human and hiPSC-CM phosphoproteomics data have been deposited to the ProteomeXchange Consortium via the PRIDE partner repository with dataset identifiers PXD057631 and PXD072789, respectively, and will be made publicly available. A subset of our human myocardial INKA kinases dataset, namely those pertaining to relevant kinases, was analyzed and published previously. This data was analyzed simultaneously as the other INKA scores reported here and, therefore, was subject to the same statistical analyses [13].

### Statistical analyses

Graphpad Prism 10 was employed for all statistical testing. Data normality was checked prior to statistical analyses. When multiple testing was applied, we corrected within tests. For all data presented, unless noted otherwise, a *P*-value of <0.05 was considered significant. A summary of all statistical tests and details for the individual analyses can be found in **Table S5**.

## Results

### Expression of enzymes involved in MT acetylation and detyrosination

Microtubular PTMs are mediated by enzymes and changes in the expression of these enzymes may explain the increased abundance of their corresponding PTMs in HCM. We therefore first studied key MT enzymes with Western immunoblots. Increased acetylation of the α-tubulin Lys40 residue in HCM compared to NF myocardium may be explained by higher αTAT1 expression and/or reduced histone deacetylase 6 (HDAC6), the enzyme responsible for MT deacetylation (**Figure 1A**). In contrast, we observed the exact opposite, with significantly lower levels of αTAT1, and higher levels of HDAC in HCM (N=10) compared to NF samples (NF, N=5). HDAC6 expression was highest in G+ patient samples (**Figures 1B**, **C**, **D** and **Figure S3A**). The increase in MT detyrosination in HCM compared to NF hearts may result from reduced tubulin tyrosine ligase (TTL) or increased levels of detyrosinating VASH1-SVBP or VASH2-SVBP complexes (**Figure 1A**). Because of their low abundance both VASH isoforms can only be detected using immunoprecipitation of VASH-SVBP complexes [32]. We were able to detect VASH1, but not VASH2, in human interventricular myocardium (**Figure 1F** and **G**, and **Figure S3B** and **S3C**). We did not observe significant changes in TTL (**Figures 1B, E**, and **Figure S3A**) or VASH1-SVBP levels in HCM (**Figures 1F** and **Figure S3B**) compared to NF. There is a trend towards increased VASH1 levels in HCM patients, which does not correlate to total detyrosination levels (**Figure S3D**, *R*^2^=0.25, *P*=0.43). Aspecific bands on the VASH membranes are likely to be heavy and light IgG chains. Overall, these data do not explain the HCM MT code.

**Figure 1.**
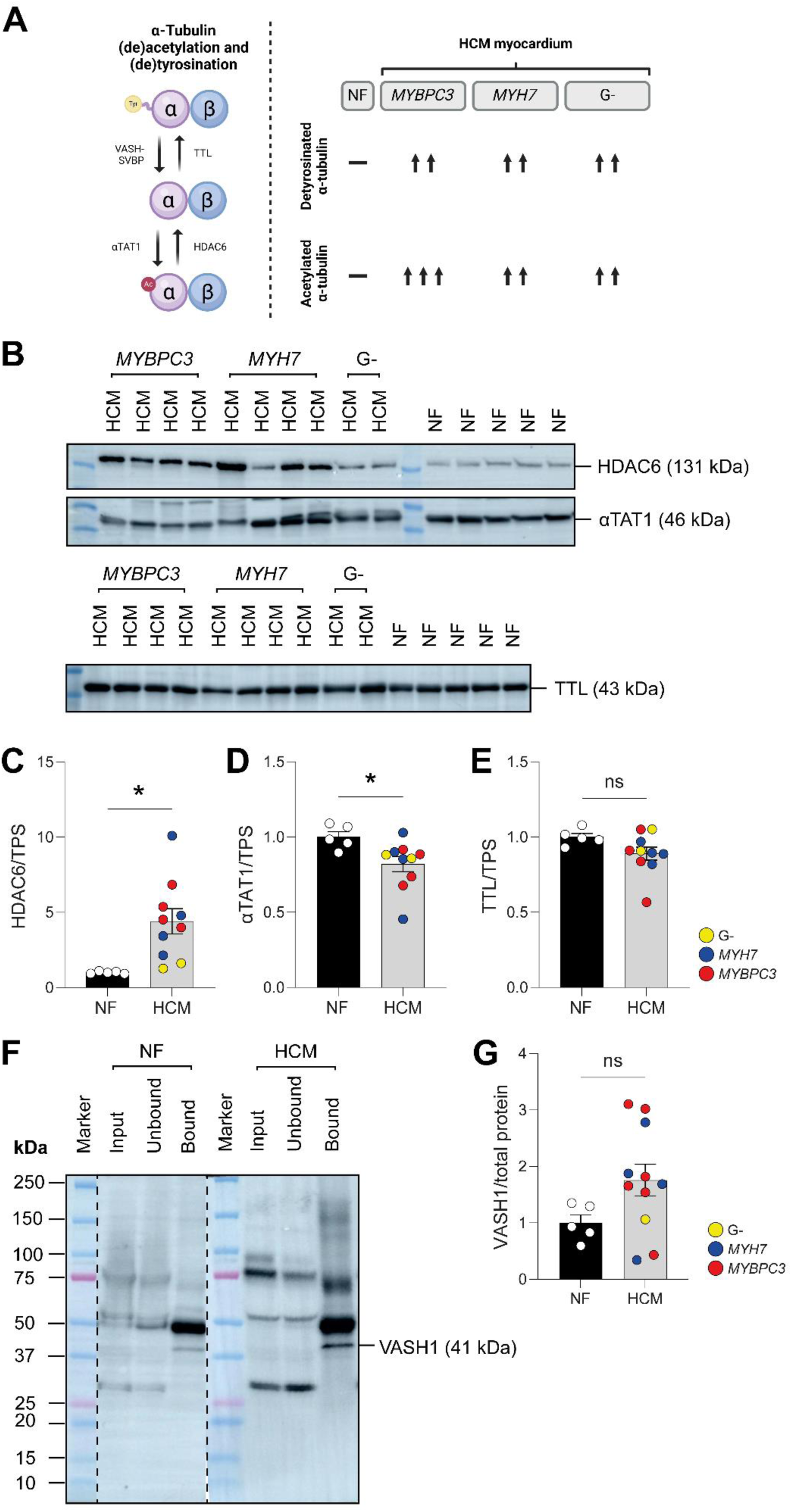
Expression of enzymes involved in post-translational modifications of microtubules in HCM. (**A**) Schematic overview showing the enzymes involved in microtubule acetylation and detyrosination. HCM patients, especially when genotype-positive, present with increased levels of total, acetylated and detyrosinated microtubules [18]. (**B**) Representative blot images. Unlabeled lanes are protein markers. (**C**-**E**) Quantified levels of HDAC6 (**C**), αTAT1 (**D**) and TTL (**E**). HCM samples of patients with different genotypes were compared to non-failing (NF) controls. (**F** and **G**) Representative blot images and quantification of VASH1 levels in HCM samples with different genotypes compared to NF samples. Every dot represents a patient or NF sample. All data was distributed normally and tested using an unpaired t-test. \**P*<0.05.

We also sought to evaluate the transcriptional levels of these enzymes. In HCM myocardium, both HDAC6 and αTAT1 transcripts remained unchanged (**Figures S3E** and **F**). As for the key detyrosinating enzymes, we found a near 50% (*P*=0.13) increase and 50% (*P*=0.08) decrease in *VASH1* and *VASH2* transcripts, respectively (**Figures S3G** and **H**). *SVBP* and *TTL* transcripts were not changed in HCM patients (**Figures S3I** and **J**).

### Proteomic and phosphoproteomic coverage in HCM

We performed phosphoproteomic profiling of cardiac tissue from 24 HCM patients and compared this to 8 NF samples. Supervised hierarchical clustering (**Figures S4B** and **S4C**) was performed as well as principal component analysis (**Figures S4D** and **S4E**) to study whether HCM and NF samples cluster separately based on their pTyr or pSer/pThr features. For the pTyr data, samples were slightly more similar to each other than for the pSer/pThr data. Despite this, distinct clusters for HCM and NF control samples were observed for both. In the pSer/pThr dendrogram two clusters can be observed (**Figure S4C**). Based on the clinical and echocardiographic parameters, these two HCM clusters show differences in the resting LV outflow tract pressure (51.1 ± 10.8 mmHg, N=12 vs 90.7 ± 11.5 mmHg, N=9, for overview **see Table S1**). We did not observe distinct phosphoproteomes when comparing genotypes, sexes or interventricular septal wall thicknesses normalized by body surface area (data not shown).

We identified 1984 pSer/pThr (*P*<0.1, **Figure 2B**), 160 pTyr sites (*P*<0.1, **Figure 2A**), and 463 proteins (*P*<0.05, **Figure 2C**) that were differentially altered in HCM compared to NF. A subset of these proteins showed both altered differential abundance and differences in their phosphorylation state, resulting in different overlapping categories (**Figure 2D**). Notably, proteins involved in the cytoskeleton (*SYNPO2L*, *SVIL*, *MAP4*, *NES*, *TUBA1B*, *SORBS2*, *DES*, *VIM*, and *ACTA1*) constitute major overlapping components (**Figure 2D**).

**Figure 2.**
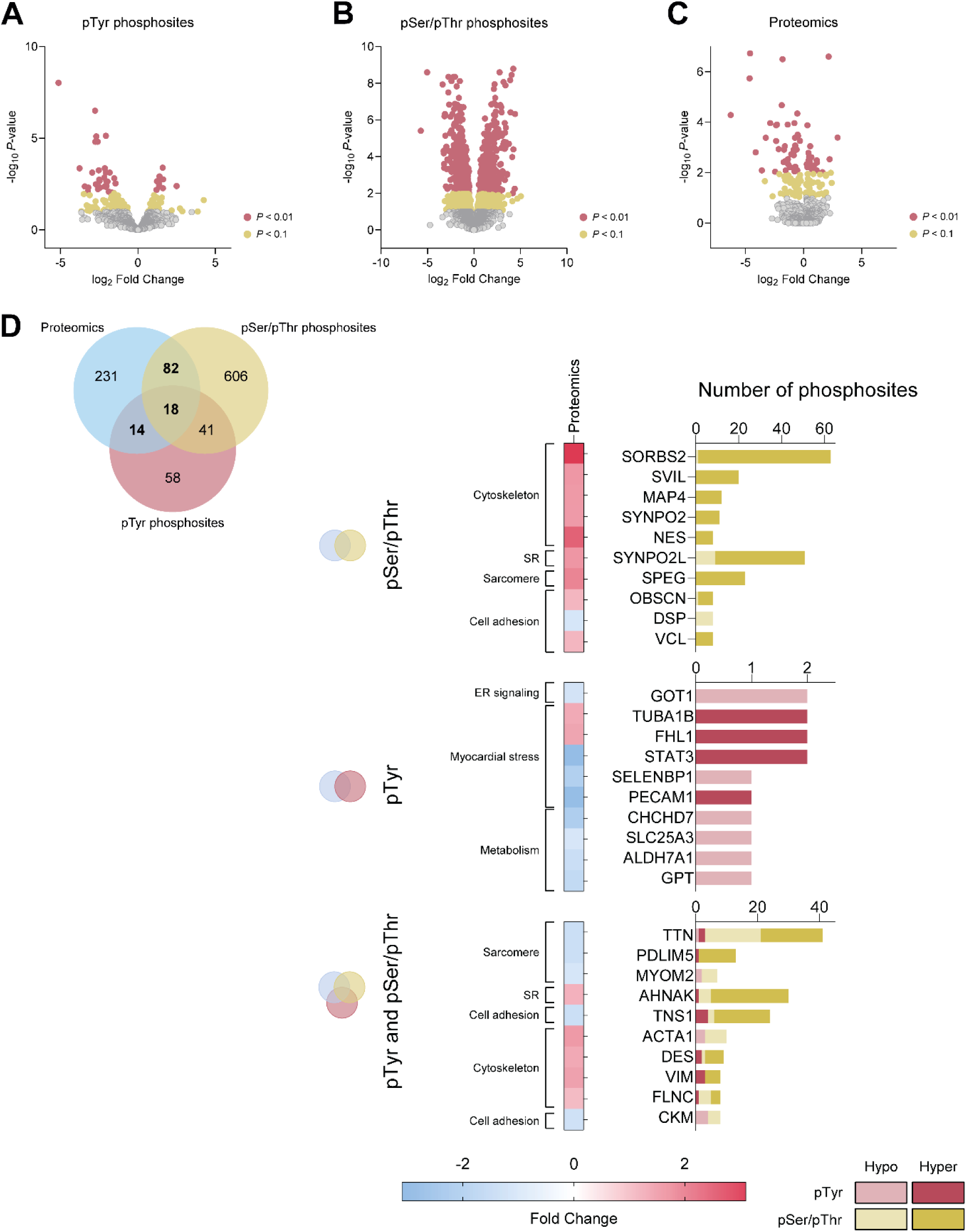
Data distribution across datasets shows diversity of phosphorylation signaling in HCM. (**A**) Volcano plot showing significantly altered pTyr sites in HCM patients (*P*-values assigned for an FDR set to 0.1). (**B**) Volcano plot showing significantly altered pSer/pThr sites in HCM patients (*P*-values assigned for an FDR set to 0.1). (**C**) Volcano plot showing significantly altered differential proteins in HCM patients compared to NF controls (*P*-values assigned for an FDR set to 0.1). (**D**) Venn diagram depicting the overlap between the differential proteomics, phosphotyrosine (pTyr) and phosphoserine and phosphothreonine (pSer/pThr) datasets based on protein hits. The top 10 hits overlapping between the differential proteomics and phosphoproteomics datasets as depicted in the Venn diagram (bold numbers) are shown in the bar charts in the right panel. Hits were selected by filtering proteins with significant changes in phosphorylation, which were first ranked by the number of differential phosphosites and then by the fold changes in their phosphosites. For every differential protein, the number of significantly altered phosphosites, either pTyr or pSer/pThr is indicated. In addition, the colours of the bars indicate what proportion of phosphosites are hypo- (lighter shades) or hyperphosphorylated (darker shades). Protein abundancy, expressed in fold changes, corresponding to these proteins are depicted on the left side of the bars.

### INKA and phosphosite-specific signature analysis

To identify which kinases are responsible for the differential phosphorylation seen in HCM tissue, we performed INKA analysis as described previously [30]. INKA allows for the quantification of kinase activity by combining kinase-centric data, i.e. the extent to which a given kinase is phosphorylated together with the phosphorylation state of substrate proteins that are phosphorylated by that kinase. Altogether, 18 Tyr kinases (**Figure 3A**) and 20 Ser/Thr kinases are significantly hyperactive in HCM, while 1 Ser/Thr kinase is hypoactive in HCM compared to NF myocardium (**Figure 3B**). The enriched Tyr kinases include EGFR, IGF1R, INSR and FGFR1, which have been linked to activation of MAPK signaling [13, 37, 38]. In line with this, several MAPKs, including MAPK1 and MAPK3 (both dual-specificity kinases), MAPK14 and MAPK10 were activated in our dataset. In addition, HCM patients displayed increased activity of the non-canonical Tyr kinases PDGFRA, FER, EPHA3, EPHA4, JAK2, CDK5, HCK, PTK2, FYN, and SRC (**Figure 3A**). Several non-canonical Ser/Thr kinases such as NEK9, DAPK3, PAK1, CSNK1E, ROCK2, MAP2K2, MARK2, BCKDK, MAP4K4, ILK and EEF2K are also more active in HCM. In addition, we found increased activation of GSK3β, PRKCE, and AMPK, which are Ser/Thr kinases that have previously been linked to hypertrophy [39] or failing myocardium of dilated cardiomyopathy patients (**Figure 3B**) [40, 41].

**Figure 3.**
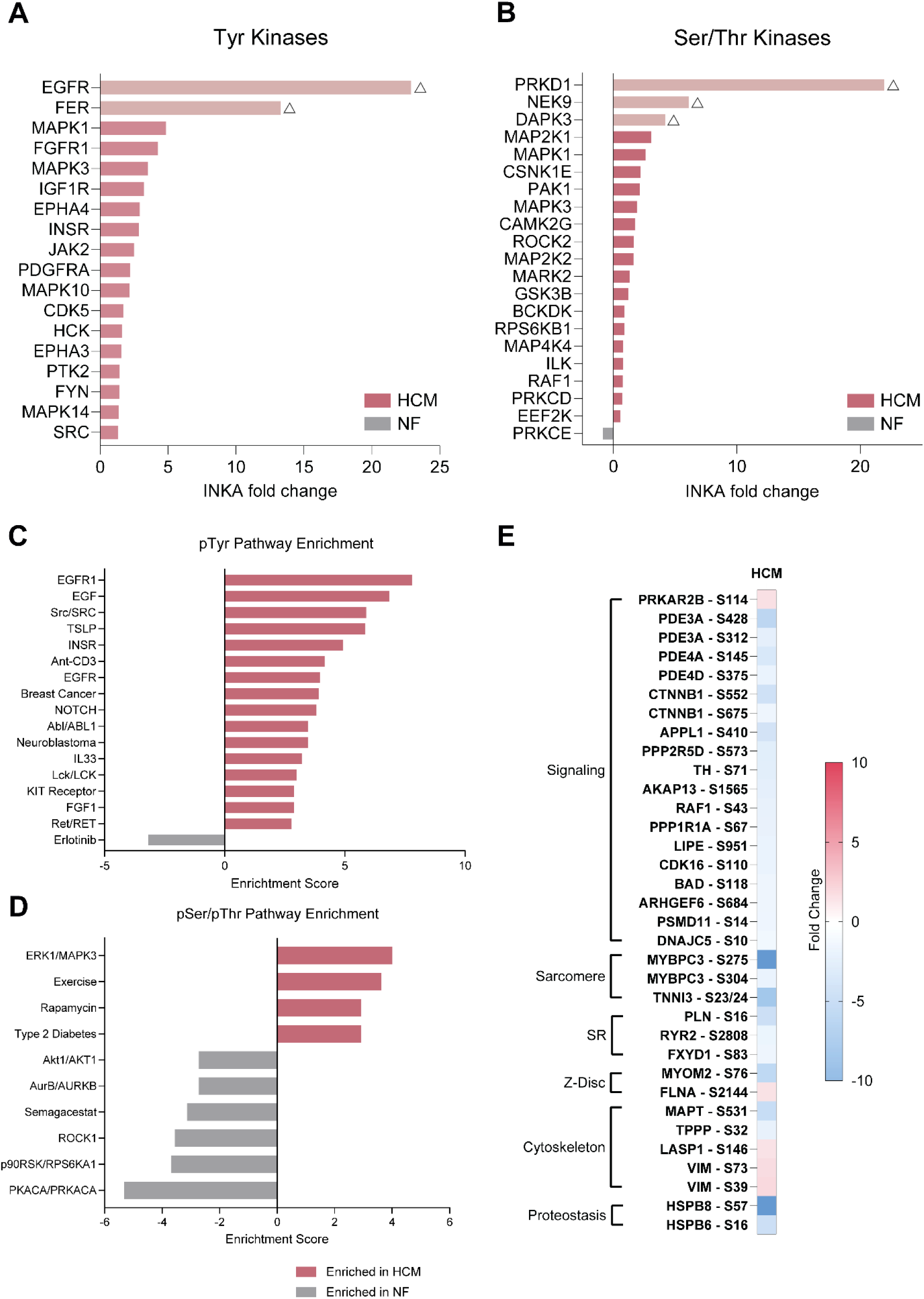
Pathway enrichment in HCM shows decreased PKA signaling. (**A**) Significantly hyperactive Tyr kinases in HCM patients when compared to non-failing donors (NF), expressed by their INKA fold changes. (**B**) Significantly hyper- and hypoactive Ser/Thr kinases in HCM patients when compared to NF, expressed by their INKA fold changes. For the INKA fold changes, normally distributed data was tested using an unpaired t-test and using Mann-Whitney if the data was not normally distributed. Lighter bars with triangles indicate kinases that had no score for NF samples, but were scored in 25% of HCM samples. Therefore, these values correspond to their delta value instead of fold change. (**C**) Phosphotyrosine (pTyr) pathway enrichment scores using the entire corresponding dataset (*P*<0.05). (**D**) Phosphoserine (pSer) and phosphothreonine (pThr) pathway enrichment scores using the entire corresponding dataset (*P*<0.05). All pathways that are enriched in HCM when compared to NF controls are indicated as pink bars, whereas those enriched in NF are depicted as gray bars. (**E**) All phosphosites corresponding to the PKACA/PRKACA, or protein kinase A (PKA) signaling pathway, which is significantly decreased in HCM patients. Fold changes show significantly (FDR<0.1) lower phosphorylation of these serine residues.

To evaluate the signaling signatures in HCM samples, we performed a phosphosite signaling enrichment analysis using PTM-SEA [31]. This analysis demonstrates an enrichment of several Tyr kinase pathways in HCM tissue, including EGFR and EGF, Src, INSR, and FGF1. As shown in **Figure 3C**, this is in line with the increased activity of these kinases. The erlotinib signaling pathway, which is an EGFR inhibitor, is also reduced in HCM compared to NF (**Figure 3C**). Significantly reduced Ser/Thr signatures in HCM compared to NF include PKACA/PRKACA, p90RSK/RPS6KA1, and ROCK1 signaling, whereas ERK1/MAPK3, exercise and rapamycin signatures are significantly enriched (*P*<0.05, **Figure 3D**). Decreased protein kinase A (PKA) signaling, a hallmark of HCM, is further exemplified by plotting the individual phosphoresidues that are included in this signature (**Figure 3E**) [11, 42]. Canonical signaling and sarcomere proteins within this signature are hypophosphorylated, whereas only a subset of phosphoresidues pertaining to the cytoskeleton are hyperphosphorylated in HCM compared to NF (FDR<0.1). Based on PhosphositePlus (v6.7.5), these sites could also be phosphorylated by PKG (S146 on LASP1), Akt1 or PKC (S39 on VIM), and PAK1 or AurB (S73 on VIM).

### Correlating phosphoresidues and the MT code

To probe which kinases are responsible for the increased levels of α-tubulin and its post-translationally modified forms in HCM patients, we set out to perform correlation analyses between INKA-based kinase scores and alterations in the MT code. Correlation analyses allow for screening of potential modulators of the altered MT code. Results were filtered for a minimum of 10 observations (N), *R*^2^≥0.3, and a *P*-value<0.05. Here, we present significant hits. Due to the low abundance of Tyr kinases, we only found weak correlations that we removed due to their low *R*^2^ values. A number of Ser and Thr kinases correlated to tubulin or tubulin PTMs. CSNK1A1 positively correlated with total, detyrosinated and acetylated α-tubulin (*R*^2^=0.33, 0.29 and 0.38, respectively, *P*<0.05). RPS6KB2 also positively correlates to both detyrosinated and acetylated α-tubulin (*R*^2^= 0.36 and 0.56, respectively, *P*<0.05), but not to total α-tubulin levels. In addition, DYRK2 positively correlates to acetylated α-tubulin (*R*^2^=0.38, *P*<0.05). Both CSNK1A1 and RPS6KB2 can be phosphorylated by PKA, but the corresponding phosphosites (S218 and S242 for CSNK1A1 and S473 for RPS6KB2) were not detected in our screen. The Y382 site on DYRK2 exhibits increased phosphorylation in HCM and can be phosphorylated by both EGFR and FGFR1 (PhosphositePlus v6.7.5), consistent with their increased activity in HCM as shown by our INKA analysis (**Figure 3A**). Although it is unknown whether Y382 induces enzymatic activity, this site is significantly more phosphorylated in our HCM samples (**Figure S4F**).

We then expanded our correlative analyses to all phosphosites. All results were filtered for a minimum of 10 observations (N), *R*^2^>0.4, and *P*-value<0.05. Here, we present the (where possible) top 10 correlations for total, detyrosinated and acetylated α-tubulin (**Table 2**). Interestingly, 3 phosphoproteins correlate to 2 or 3 of the microtubular markers. S951 on hormone-sensitive lipase (LIPE) negatively correlates to total (*R*^2^=0.763, *P*<0.0001) and detyrosinated tubulin (*R*^2^=0.590, *P*= 0.0002). S1949 on MAP1B and S286 on cMyBP-C both show significant correlations to detyrosinated (*R*^2^=0.609 and 0.561, respectively) and acetylated α-tubulin (*R*^2^=0.605 and 0.530, respectively). HCM patients carrying a mutation in the *MYBPC3* gene typically present with the highest increase in acetylated α-tubulin in conjunction with reduced expression of cMyBP-C [18]. With the exception of acetylated α-tubulin, we were unable to observe clustering of specific sarcomere genotypes and instead found that G- patients tended to cluster together for correlations between cMyBP-C S286 and the MT code (**Figures S4G-I**). As this phosphoresidue is located in the M-domain of cMyBP-C, we also correlated other well-studied phosphoresidues within this protein, and found that with the exception of S284 and S311, all of them significantly correlated with alterations in the MT code (**Figure S4J**).

**Table 2.**
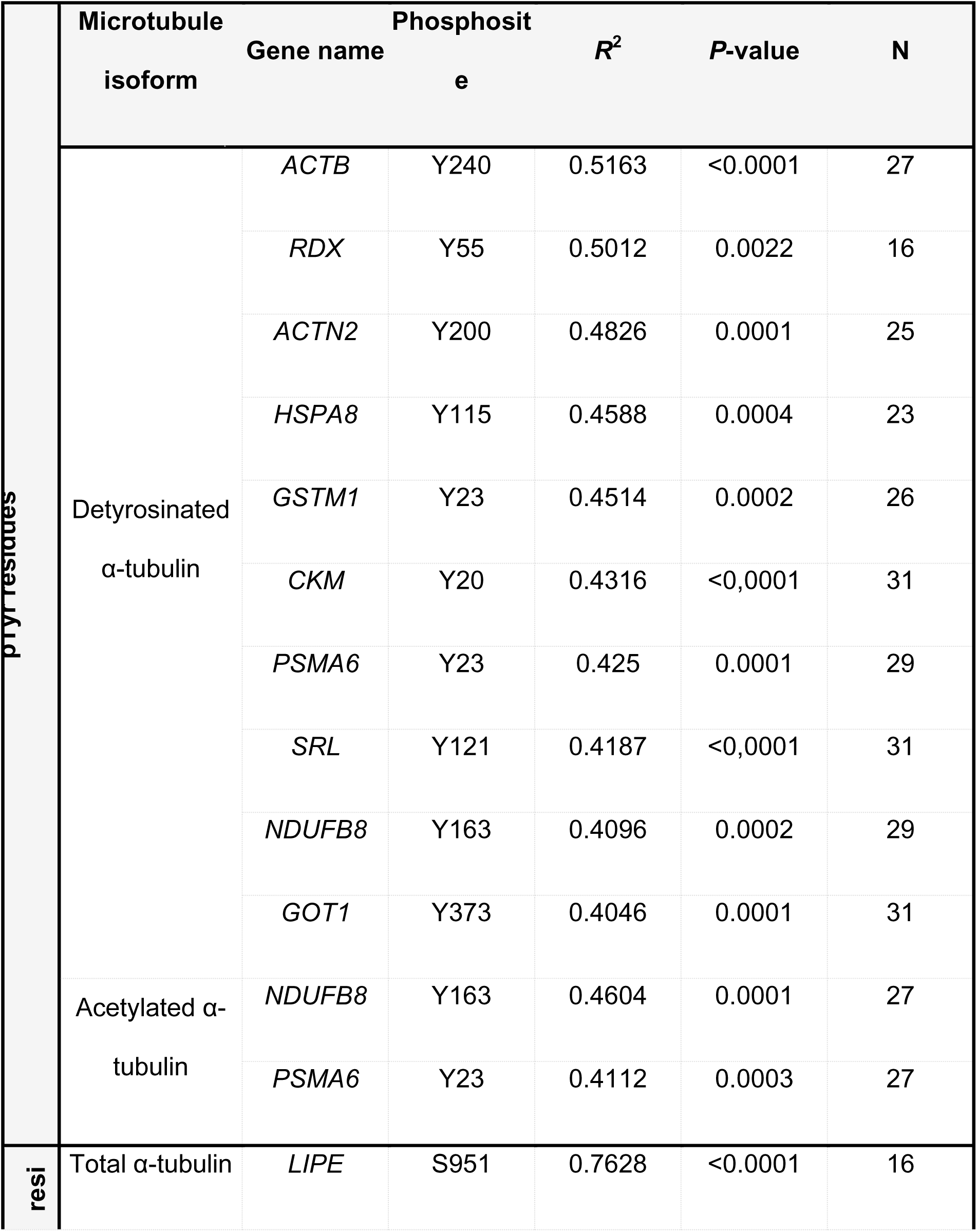
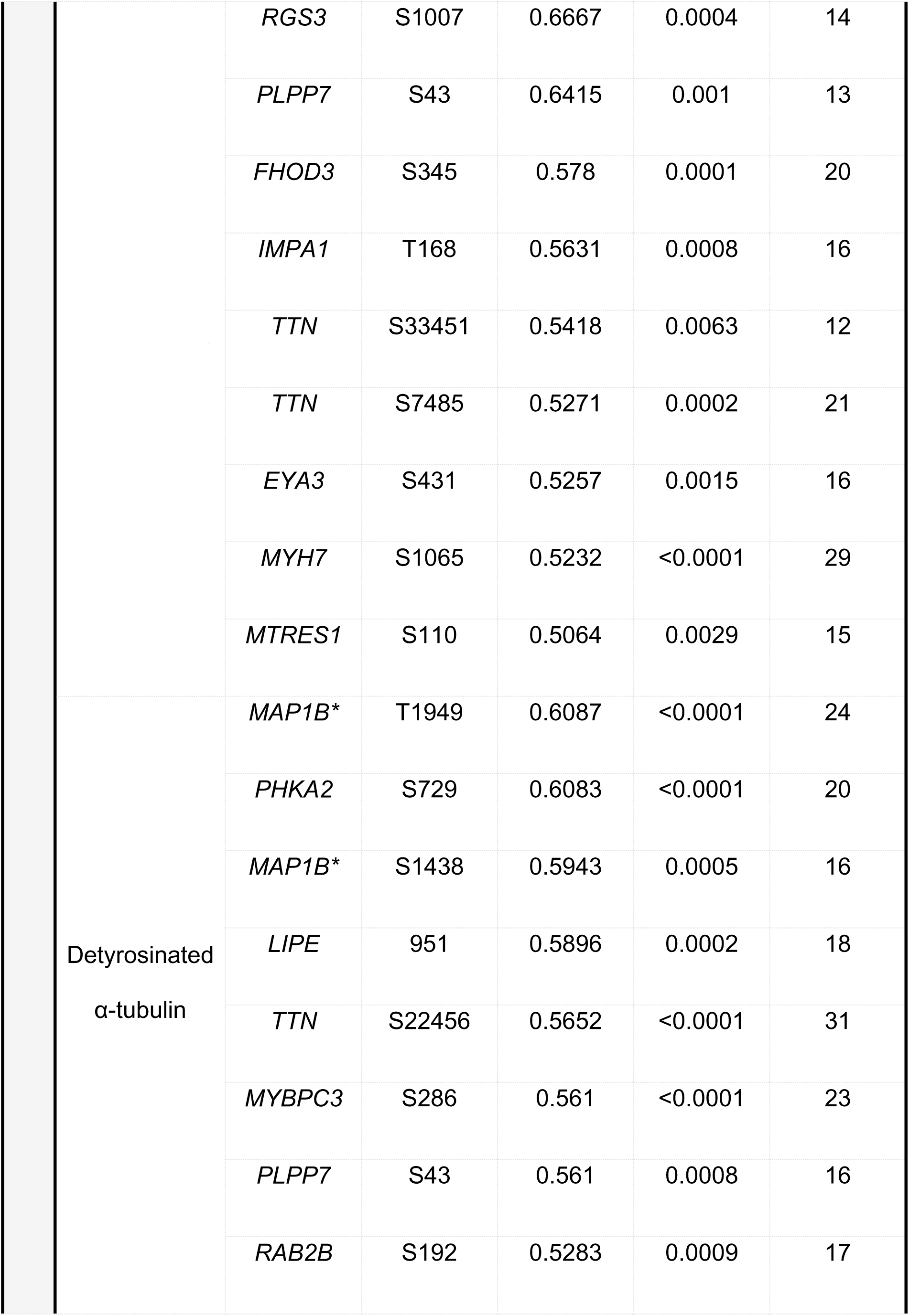
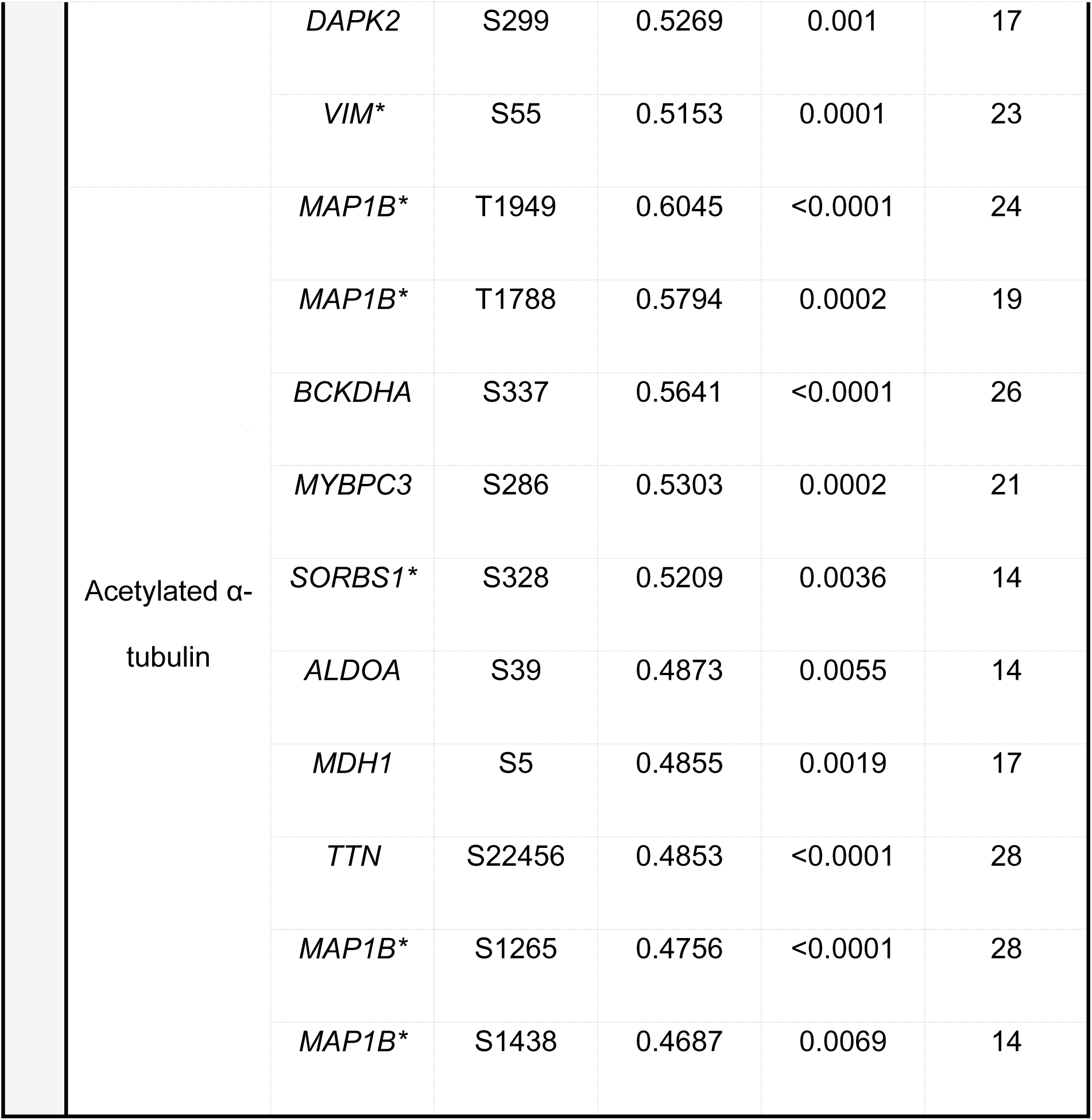
Top correlations for phosphotyrosine residues (top, pTyr), phosphoserine/phosphothreonine (bottom, pSer/Thr) and microtubule codes in HCM and NF samples. *Positive correlation.

PhosphositePlus (v6.7.5) did not report any upstream kinases for the pTyr residues that we found. S951 on LIPE and S275 and S304 on cMyBP-C are all phosphorylated by PKA. Though upstream kinases of the phosphosites that we found on MAP1B have not been reported yet, these sites have been linked to Aurora B kinase inhibitor, EGF and ischemia-driven pathways. With the exception of S337 on BCKDHA, which can be phosphorylated by BCKDK and is significantly more active in HCM, no upstream kinases have been linked to the remaining pSer/pThr residues that significantly correlate to the altered MT code.

### Link between phosphokinome and cardiac pathology

To determine if we could link specific kinases to cardiac hallmarks of disease, we correlated our clinical data to the INKA, and subsequently to our whole phosphoproteomics dataset. All HCM patients showed impaired diastolic function, as is indicated by increased LV filling pressures (E/e′) and atrial dilation (LADi), and interventricular septal hypertrophy (IVSi) (**Table 1**). Here, too, we correlated kinase activity (INKA) data with clinical parameters of diastolic dysfunction and present only significant hits. We found that the Tyr kinase HIPK1 positively correlates with E/e’ (*R*^2^=0.35, *P*<0.01) and that EPHA4 and EPHA5 also positively correlate with LADi (*R*^2^=0.44 and 0.44, respectively, *P*<0.05). EPHA5 can be phosphorylated by FGFR1 on Y833, a phosphosites that is significantly increased in our HCM samples. Another member of the EphA kinase family, Tyr kinase EPHA2 positively correlates with the resting LV outflow tract gradient (LVOTg (*R*^2^=0.33, *P*<0.01). Phosphorylation on Y544 on EPHA2 indicates enzymatic activity and shows a trend, though statistically insignificant, towards an increase in our HCM samples. Overall, little has been reported on these kinases and their activation.

Ser/Thr kinases correlated to LVOTg. PRKD2, PRKACG, and PRKACB all positively correlate with LVOTg (*R*^2^=0.31, 0.36, and 0.46, respectively, *P*<0.05). PRKCI, BRAF and MARK1 all negatively correlate with LADi (*R*^2^=0.41, 0.44, and 0.37, respectively, *P*<0.05). Thus, when integrating phosphoproteomic and clinical data, in addition to known signaling pathways, e.g. reduced PKA signaling, kinases that previously have not been linked with HCM, such as those in the EphA family, are identified and may underlie cardiac dysfunction and remodeling.

Next, we correlated all significantly altered phosphosites with clinical parameters. Again, by ranking these hits according to their *R*^2^ values and filtering for a *P*-value<0.05 and *R*^2^>0.4, we identified candidates that cover different functions within the cardiomyocyte (**Table 3**). Diastolic dysfunction characterized by e’, E/e’, and LADi, significantly correlates with T284 in *EHBP1L1* (*R*^2^=0.560, *P*<0.05), S646 on *CLASP1* (*R*^2^=0.593, *P*<0.05), and S14 on *SORBS2* (*R*^2^=0.808, *P*<0.05), respectively. LV hypertrophy, i.e. IVSi, significantly correlates with S149 on *TPD52L1* (*R*^2^=0.660, *P*<0.05). The LVOTg correlates with T649 on *CTNNA3* (*R*^2^=0.621, *P*<0.05). By using PhosphositePlus (v6.7.5), we sought to identify upstream kinases and/or regulatory kinases of the phosphosites we report in **Table 3**. For the pTyr residues, we only found that Y217 is phosphorylated by Abl1. For the pSer/pThr residues, we found that S895 on LMO7 and S1757 on NUMA1 can be phosphorylated by PRP4, T4564 on AHNAK by TBK1, and S608 on STIM1 by ERK2. We, therefore, were unable to link the phosphosites to common upstream kinases.

**Table 3.**
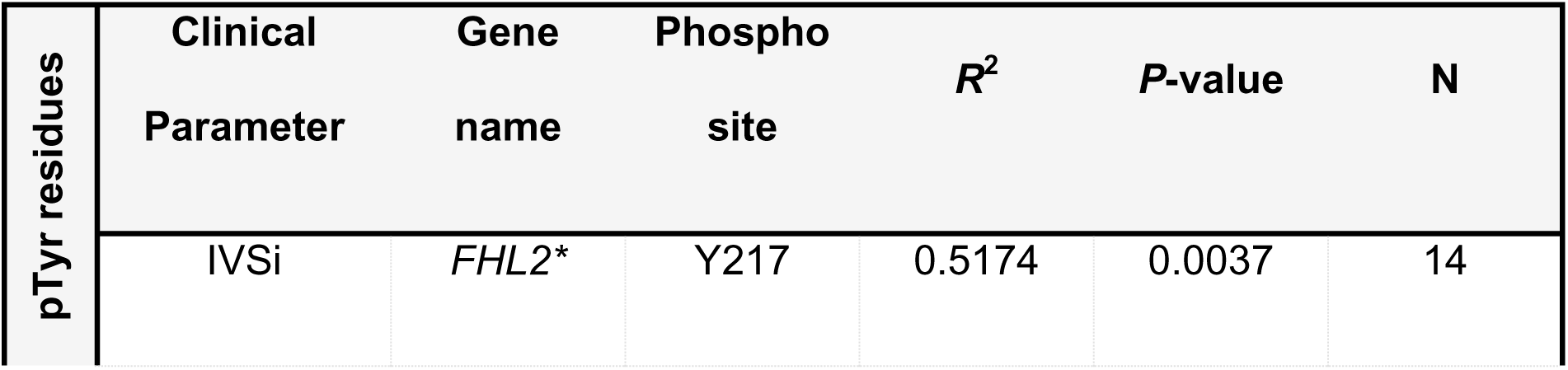
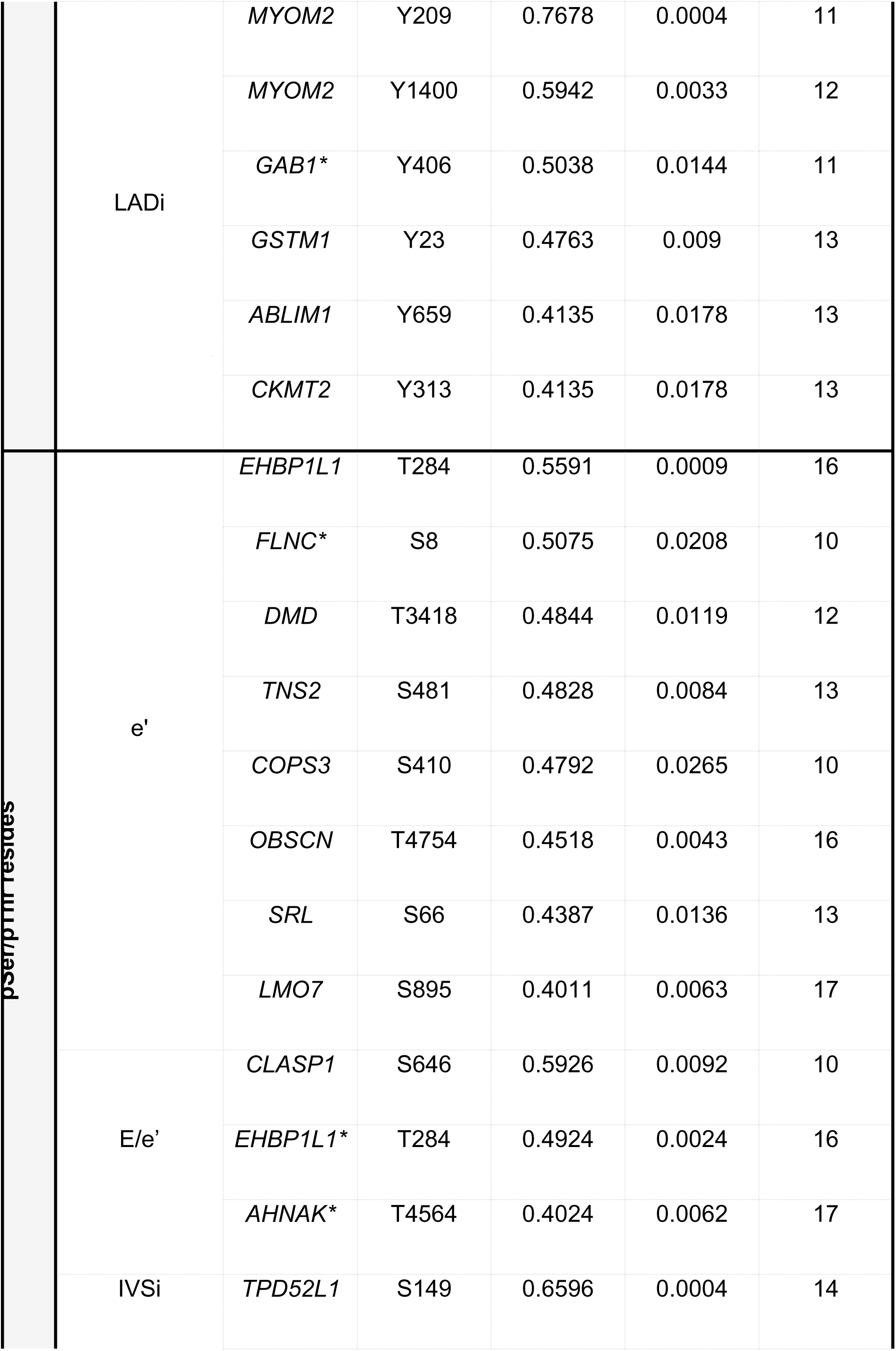
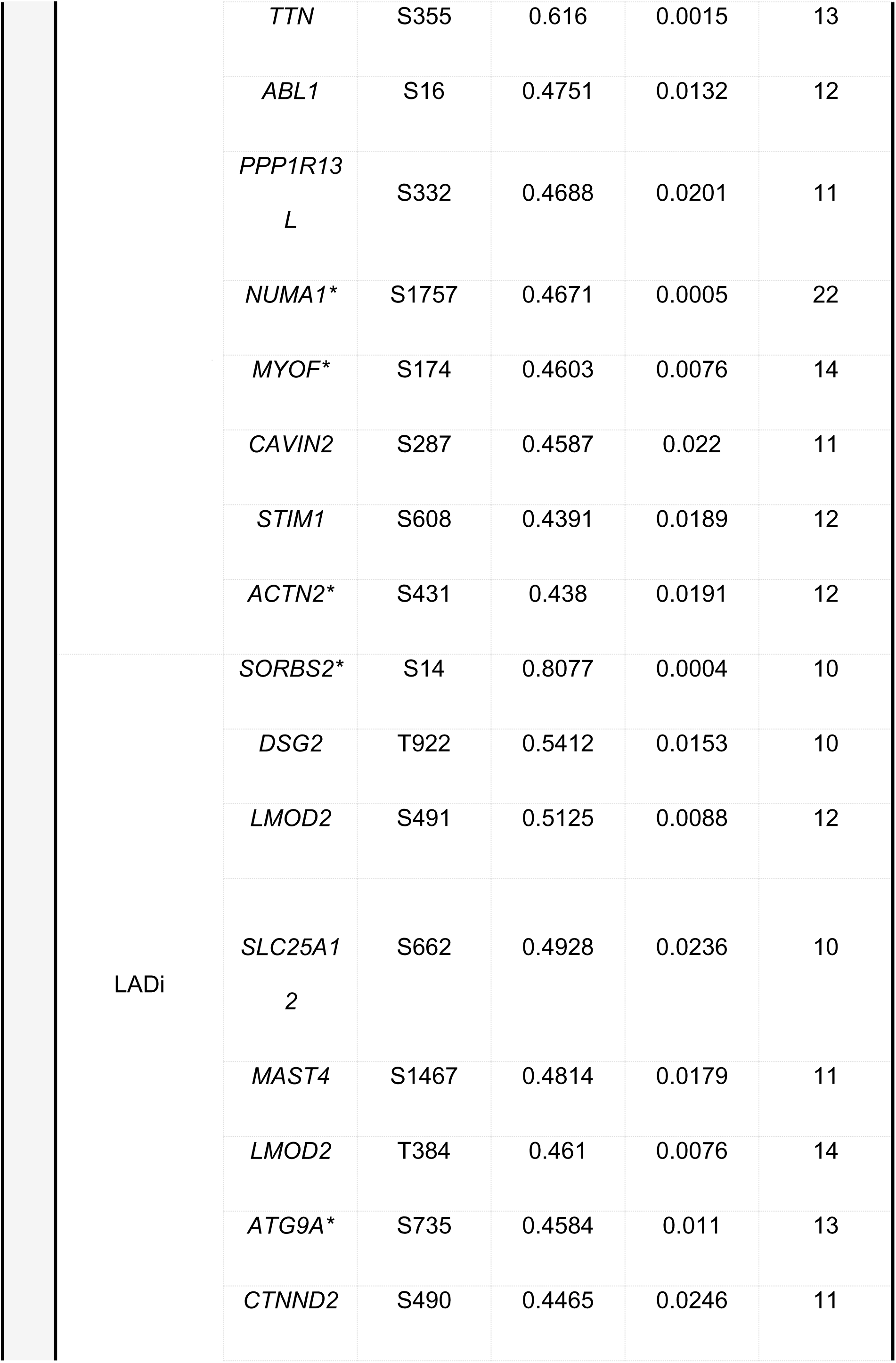
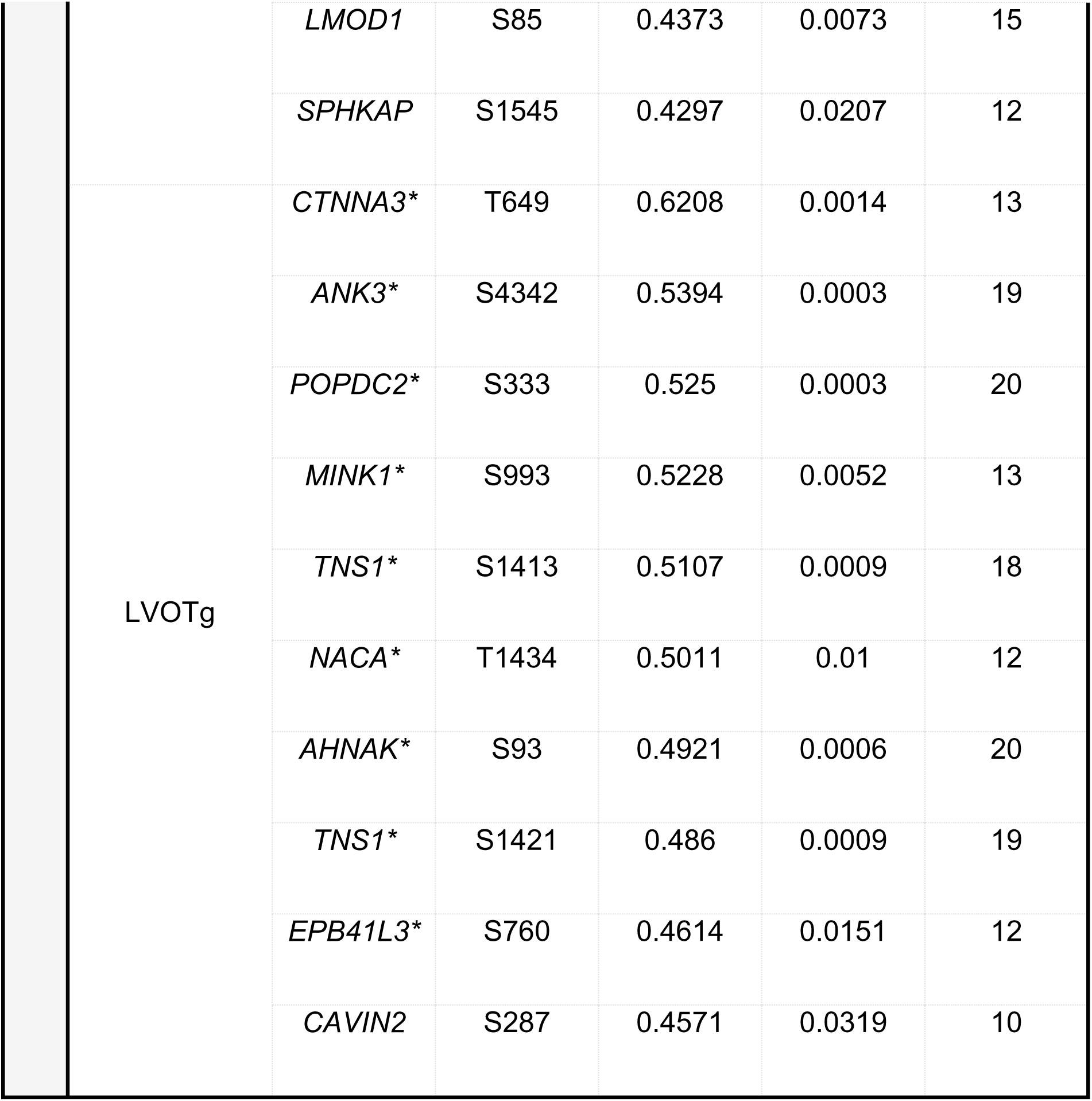
Top correlations for phosphotyrosine residues (top, pTyr), phosphoserine/phosphothreonine (bottom, pSer/Thr) and clinical parameters in HCM patients. *Positive orrelation.

### HOM_Arg943X_ hiPSC-CMs exhibited an increased MT repolymerization at 60 days after start of differentiation, which was destabilized through PKA activation

Next, we sought to evaluate the etiological position of MT changes in HCM pathology. To this end, we utilized an hiPSC line with mTag-RFP-T-TUBA1B to introduce a Dutch founder mutation for HCM (Arg943X) using CRISPR/Cas9 genetic tools. Protein analyses of hiPSC-CMs showed that HOM hiPSC-CMs are deficient of cMyBP-C and both showed similar levels of slow skeletal troponin I and phospholamban (PLN) (**Figure S5**). RT-qPCR of a sarcomeric maturity panel confirmed that at 30 days after start of differentiation, both WT and HOM hiPSC-CM lines did not show significant differences (**Figure S6**).

To first define the impact of the homozygous Arg943X mutation on hiPSC-CM contractility, we utilized a high-throughput and optical method to simultaneously measure contractility and Ca^2+^-transients in 30-days old hiPSC-CMs. Compared to their WT counterparts (0.57 Hz) depicted in **Figure S5E**, unpaced HOM_Arg943X_ hiPSC-CMs showed a slightly decreased baseline beating frequency (0.56 Hz, *P*<0.05). When paced at 1 Hz, normalized trace averages of both pixel and calcium can be approximated for both WT and HOM lines as depicted in **Figures 4A** and **4B**. Paced HOM_Arg943X_ hiPSC-CMs presented with hypercontractility, an unaltered relaxation time, and an increased relaxation time over contraction time ratio (RT/CT) of HOM_Arg943X_ hiPSC-CMs (**Figures 4C**-**F**). In line with this, we observed a decreased calcium release time, an unaltered calcium reuptake time, and a shortened calcium peak width time in HOM_Arg943X_ hiPSC-CMs (**Figures 4G**-**I**). However, the normalized calcium peak height of HOM_Arg943X_ hiPSC-CMs, as well as their normalized calcium departure and return velocities were decreased (**Figures 4J**-**L**). Strikingly, at 60 days after differentiation, we did not observe differences in the contraction or relaxation times, peak width or RT/CT of HOM_Arg943X_ compared to WT hiPSC-CMs (**Figures S7C**-**F**). Instead, they showed an increased calcium release time and peak width (**Figures S7G**-**I)**. We also identified a severely slowed calcium handling profile in HOM_Arg943X_ hiPSC-CMs, as reflected by their decreased calcium departure velocity, return velocity and peak (**Figures S7J**-**L**). Thus, the HCM Dutch founder mutation Arg943X induced both a hypercontractile phenotype and slower calcium handling in HOM_Arg943X_ hiPSC-CMs.

**Figure 4.**
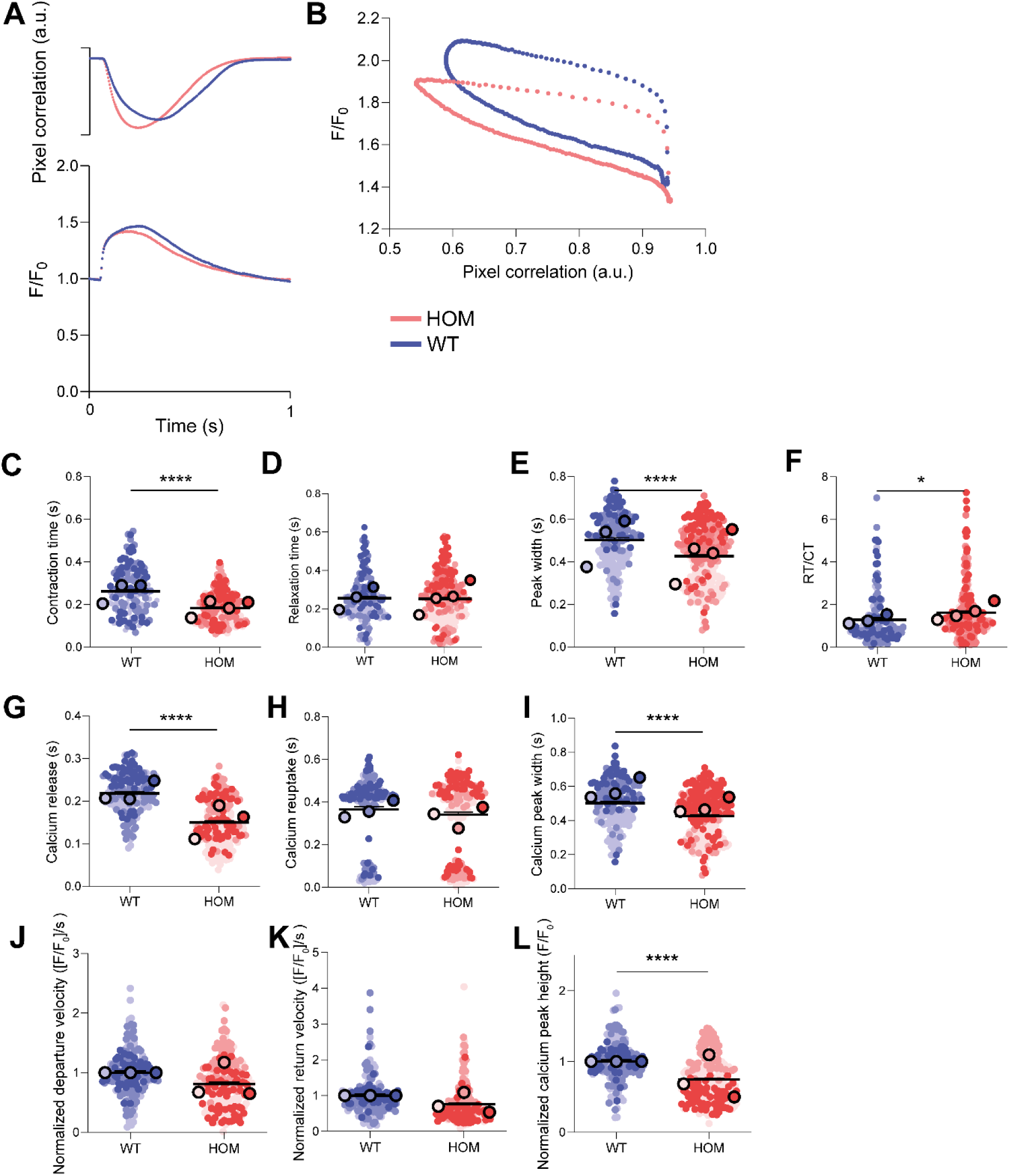
30-days old homozygous (HOM_Arg943X_) hiPSC-CMs present with an HCM-like hypercontractile phenotype. (**A**) Normalized averages of individual contractility and calcium traces obtained using 1 Hz pacing. All contractility and calcium data was tested using hierarchal cluster analysis. (**B**) Pixel-correlation and calcium loops which show how the calcium trace correlates to the pixel deviations (i.e. approximated contraction curves). When paced at 1 Hz, these cells displayed a decreased contraction time (*P*<0.001, **C**), unaltered relaxation time (**D**), decreased peak width (*P*<0.001, **E**), and an increased relaxation time over contraction time (RT/CT) ratio (*P*<0.05, **F**). These data align with a decreased calcium release time (*P*<0.001, **G**), unchanged calcium reuptake time (**H**), and decreased calcium peak width (*P*<0.001, **I**). Nevertheless, compared to WTs, HOM_Arg943X_ hiPSC-CMs showed a trend towards a decreased normalized calcium departure velocity (**J**), decreased normalized calcium return velocity (**K**) and a decreased normalized peak height (*P*<0.001, **L**). Normalized values are presented to account for different periods during which the measurements took place. Here, too, every larger dot represents an independent differentiation, with colors showing matching differentiations. Smaller dots represent individual cardiomyocytes that were measured across at least 3 wells per differentiation batch and are color-matched with the larger dots.

The age or, rather, disease stage at which a sarcomere protein alteration leads to a densified MT population in cardiomyocytes is unresolved. In our HOM_Arg943X_ and WT hiPSC-CMs, we observed α-tubulin (**Figures 5A**, **B** and **Figure S8A**), detyrosinated α-tubulin (**Figures 5A**, **C** and **Figure S8A**) and acetylated α-tubulin between 14 and 60 days after start of differentiation (**Figures 5A**, **D** and **Figure S8A**). Overall, levels of these MT PTMs were not consistently increased in HOM_Arg943X_ hiPSC-CMs. Interestingly, however, 30-days old HOM_Arg943X_ hiPSC-CMs already exhibited a trend towards increased levels of *HDAC6* (**Figure S8E**) and *VASH1* (**Figure S8F**) transcripts, albeit not significant.

**Figure 5.**
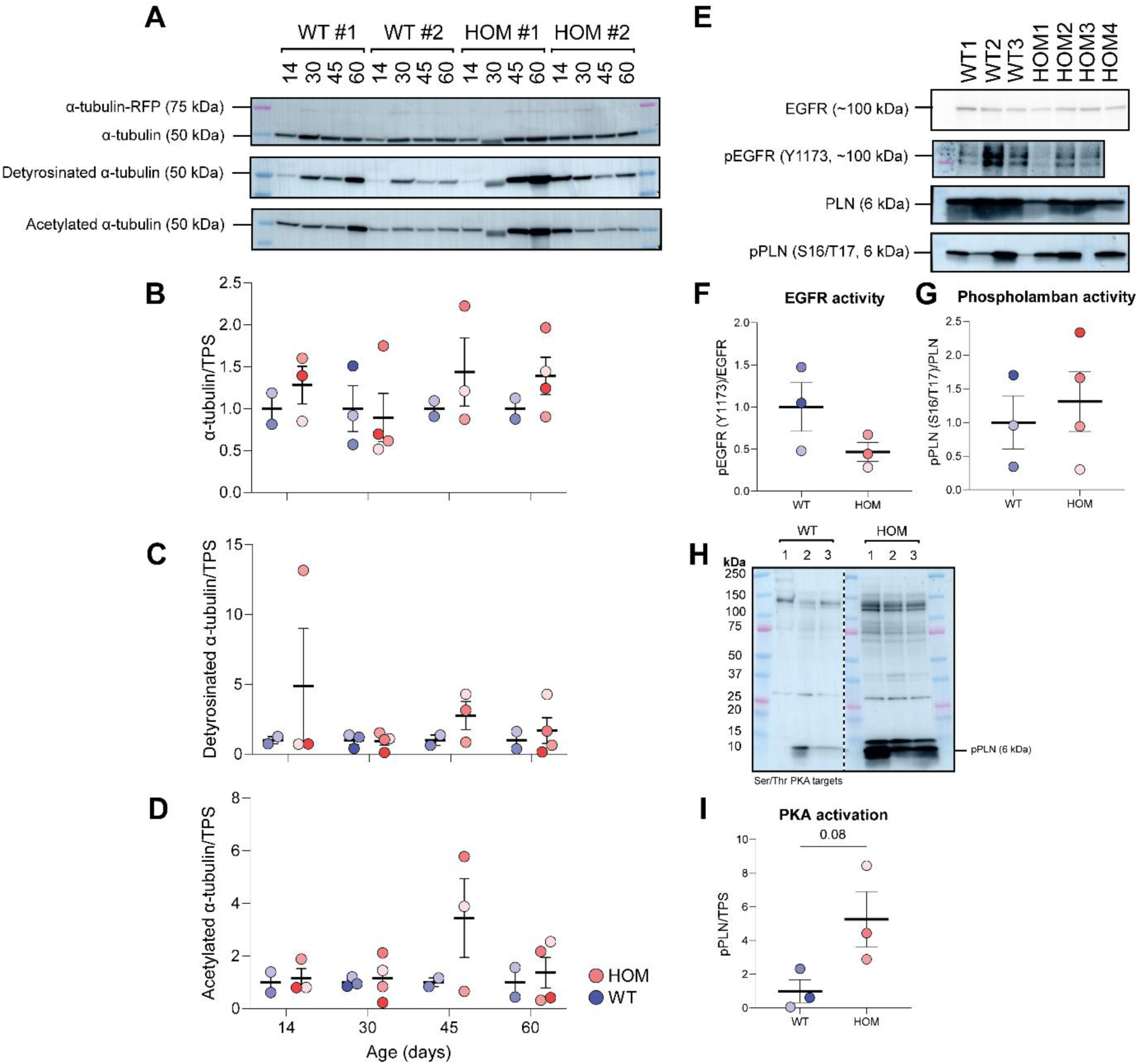
The microtubule code and phosphorylation of epidermal growth factor (EGFR) and protein kinase A (PKA) target proteins in wild-type (WT) and homozygous (HOM_Arg943X_) hiPSC-CMs. (**A**) Representative blot images of corresponding detections of α-tubulin, detyrosinated α-tubulin, and acetylated α-tubulin using WT and HOM_Arg943X_ hiPSC-CMs. The numbers indicate the ages of the samples at the time of collection. HOM_Arg943X_ hiPSC-CMs do not necessarily express significantly more α-tubulin (**B**) or its post-translationally modified forms (**C** and **D**) between 14 and 60 days after start of differentiation. In all graphs, every dot represents an independent differentiation, with colors showing matching differentiations within experiments, not in between. Smaller grey dots indicate technical replicates (i.e. images). All protein data was either tested using a two-way ANOVA or with an unpaired t-test. (**E**) Representative blot images of the detections of total EGFR, phosphorylated EGFR (Y1173 site), native phospholamban (PLN), and phosphorylated PLN (S16/T17 sites). 30-days old HOM_Arg943X_ hiPSC-CMs display a trend towards lower pEGFR (**F**) and increased pPLN (**G**) levels. Because PLN is not exclusively phosphorylated by PKA, we also looked at the detection of overall pSer/pThr PKA targets. (**H**) Representative blot images of the detection of such PKA substrates. (**I**) 30-days old HOM_Arg943X_ hiPSC-CMs presented with increased levels of PKA-specific pPLN.

Since our novel hiPSC-CM model recapitulates the HCM phenotype, we employed this HCM model to dissect whether an altered kinase activity could onset changes in the MT code. We previously highlighted two kinases, EGFR and PKA, which are hyper- and hypoactive in HCM myocardial tissue, respectively, by linking and annotating them to MT code changes using correlative analyses and PhosphositePlus v6.7.5. Therefore, we next sought to analyze baseline activity levels of EGFR and PKA in our HOM_Arg943X_ hiPSC-CMs. In contrast to our observations in HCM myocardium, the 30-days old HOM_Arg943X_ hiPSC-CMs presented with a trend towards decreased EGFR activity (**Figures 5E**, **F** and **Figures S8B** and **C**). Moreover, the HOM_Arg943X_ hiPSC-CMs displayed a trend in increased phosphorylation of Ser16/Thr17 on PLN, a downstream target of PKA. However, because PLN is not exclusively phosphorylated by PKA, we also assessed overall pSer/pThr substrate phosphorylation downstream of PKA (**Figure 5H**). Again, we found that our HOM_Arg943X_ hiPSC-CMs tended to present with increased PKA-mediated PLN phosphorylation compared to WT (**Figure 5I**).

Importantly, whole cell protein analysis as performed above decreases the spatial resolution, especially because MT populations could differ between hiPSC-CMs within one biological replicate. Moreover, such analyses provide little information as to whether MTs are more long-lived and therefore more stable. To resolve this, we performed live cell imaging experiments by incubating both WT and HOM_Arg943X_ hiPSC-CMs with the MT depolymerizing agent nocodazole at both 30 (**Figure S9** and **movies S1**-**S4**) and 60 days after the start of differentiation (**Figure 6** and **Figure S10**). In doing so, we were able to quantify the rate with which MTs depolymerize during nocodazole treatment (**Figure 6A**, panel 2) compared to baseline (**Figure 6A**, panel 1). By washing out the nocodazole, we were also able to quantify the rate with which MT repolymerization occurred (**Figure 6A**, panels 3 and 4). Using the depolymerization curve fitted in **Figure S9B**, we found that at baseline, 30-days old HOM_Arg943X_ hiPSC-CMs showed a higher percentage of α-tubulin-mTag-RFP (**Figure S9D**, *P*<0.001). Notably, during repolymerization, we found that the MTs in 30-days old HOM_Arg943X_ hiPSC-CMs recovered more quickly after the first washout as indicated by the increased AUC (**Figure S9L**, *P*<0.01). At 60-days after differentiation, HOM_Arg943X_ hiPSC-CMs showed consistently increased AUCs at baseline, after depolymerization and also after both washouts (**Figures 6D-G**), although at baseline HOM_Arg943X_ hiPSC-CMs displayed similar α-tubulin-mTag-RFP percentages as present in the WTs (**Figure S10**). Instead, at this age, we observed an overall shift towards increased MT repolymerization. Overall, our data suggested a more pathological repolymerization of HOM_Arg943X_ hiPSC-CM MTs at a later age.

**Figure 6.**
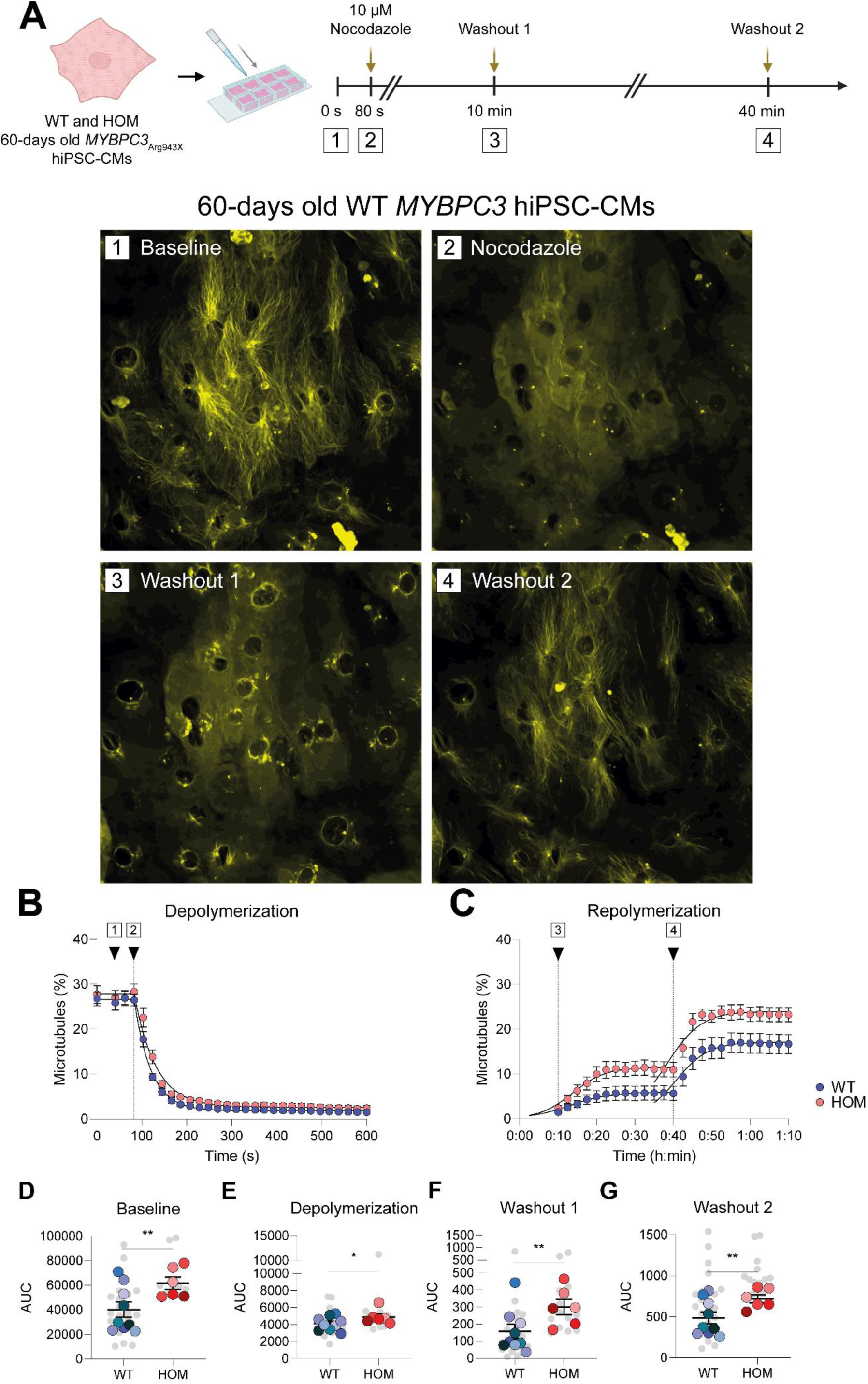
Homozygous (HOM_Arg943X_) hiPSC-CMs have more stable microtubules, which are destabilized upon isoprenaline (ISO) treatment. (**A**) 60-days old WT and HOM_Arg943X_ hiPSC-CMs were plated onto an 8-well ibidi at least 5 days prior to an experiment. The lower panel depicts representative images of both genotypes at baseline (1), after nocodazole treatment (2), after the first washout (3) and after the second washout (4). (**B**) Depolymerization trace depicting the percentage of microtubules in the cells at baseline and during/after the nocodazole treatment. (**C**) Repolymerization trace depicting the percentage of microtubules in the cells during both washouts. The area under the curve (AUC) was calculated for the baseline (**D**), depolymerization (**E**), washout 1 (**F**) and washout 2 (**G**) graphs.

To further evaluate the functional consequences of EGFR and PKA, we incubated 60-days old WT hiPSC-CMs with an EGFR inhibitor (EGFRi) or isoprenaline (ISO), which activates PKA, and performed live cell imaging (**Figures S11** and **S12**, respectively). Intriguingly, EGFR inhibition appeared to drive an increased MT repolymerization (**Figure S11C**). This increase was not in line with increased EGFR activity in the presence of MT densification in HCM patient myocardium. Instead, when we treated 60-days old HOM_Arg943X_ hiPSC-CMs with ISO (**Figures 7A**-**C**), we found decreased MT repolymerization as indicated by consistent decreases of the AUC at both baseline and during repolymerization (**Figures 7D**-**G**) as well as other parameters (**Figure S12**). In doing so, we demonstrated that PKA activation through ISO treatment mediates a decrease in MT repolymerization in HOM_Arg943X_ hiPSC-CMs. Thus, we showed that activated beta-adrenergic signaling suppresses the stabilization of MTs in HCM hiPSC-CMs.

**Figure 7.**
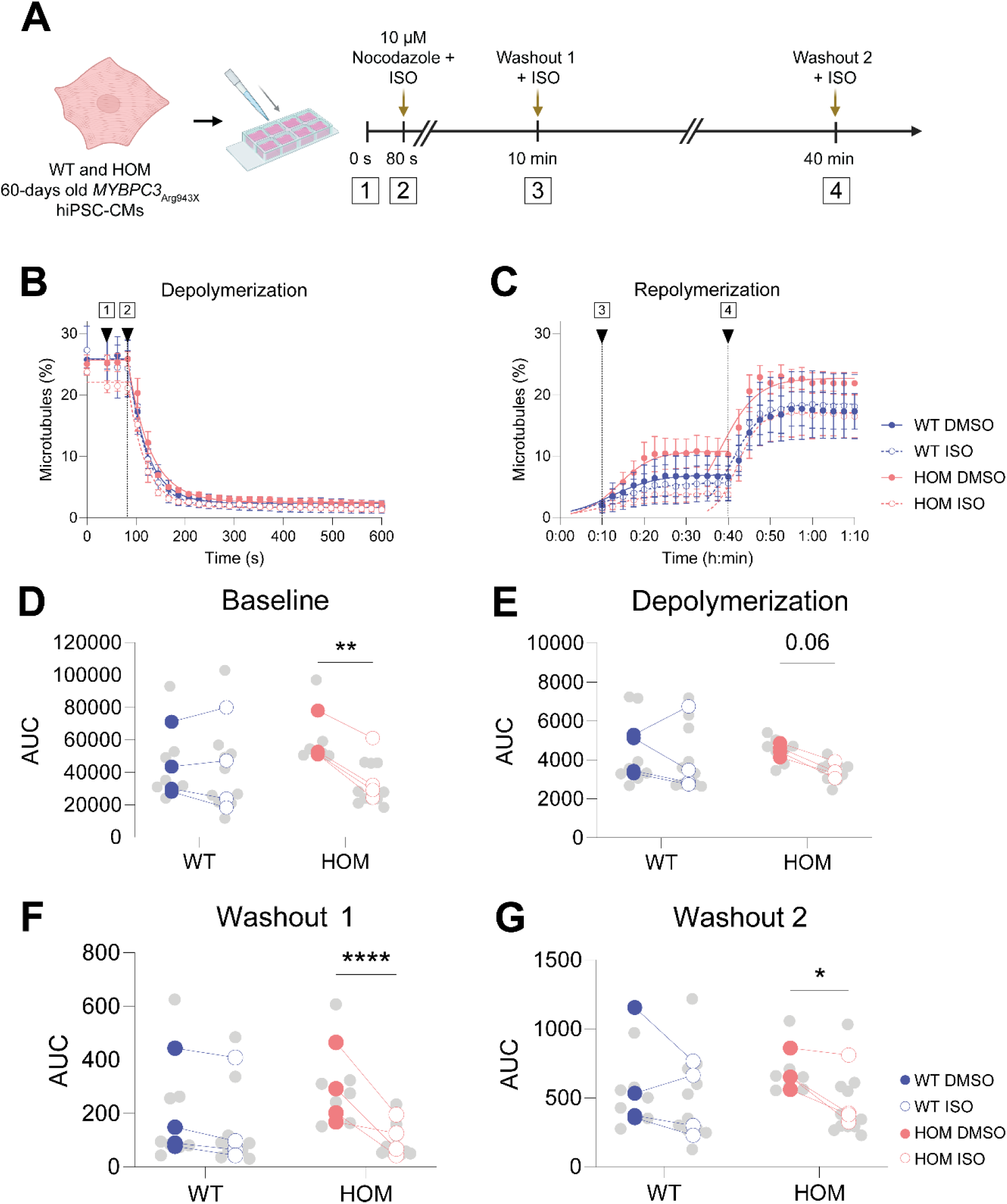
The increased microtubule repolymerization of homozygous (HOM_Arg943X_) hiPSC-CMs is destabilized upon isoprenaline (ISO) treatment. (**A**) 60-days old WT and HOM_Arg943X_ hiPSC-CMs were plated onto an 8-well ibidi at least 5 days prior to an experiment. Live cell imaging experiments were then carried out in the presence of ISO to check for any microtubule-related effects. (**B**) Depolymerization trace depicting the percentage of microtubules in the cells at baseline and during/after the nocodazole + ISO treatment. (**C**) Repolymerization trace depicting the percentage of microtubules in the cells during both subsequent washouts. The AUC values for the ISO-treated hiPSC-CMs are depicted at baseline (**D**), depolymerization (**E**), washout 1 (**F**) and washout 2 (**G**) graphs. In all graphs, every dot represents an independent differentiation, with colors showing matching differentiations within experiments, not in between. Smaller grey dots indicate technical replicates (i.e. images). All live cell data was tested using hierarchal cluster analyses.

### Inhibiting MT detyrosination decreased MT repolymerization in HOM_Arg943X_ hiPSC-CMs

Although HOM_Arg943X_ hiPSC-CMs did not show a consistent increase in MT detyrosination and acetylation compared to WT, both PTMs could help maintain the long-lived stability of MTs. To address such functional consequences, we performed live cell imaging using the MT detyrosination inhibitor epoY (**Figures S13A-N**) and the HDAC6 inhibitor tubacin, which increases MT acetylation (**Figures S13O-AA**). Our data demonstrated that in 60-days old WT hiPSC-CMs epoY, but not tubacin, decreased the AUC at baseline and after the washouts (**Figures S13K** and **M**). We, therefore, also incubated 60-days old HOM_Arg943X_ hiPSC-CMs with epoY. EpoY mediated a decrease in the MT percentage after depolymerization and the washouts (**S13E**-**G**), in addition to decreasing the AUCs at baseline and after the washouts (**Figures S13M**-**N**). These interventions thus showed that inhibiting MT detyrosination corrects the increased MT repolymerization observed in the HCM hiPSC-CM line. Overall, we demonstrate that modulating detyrosination, but not acetylation, alters MT repolymerization in both WT and HOM hiPSC-CMs.

### Validating the role of PKA in modulating microtubules

We validated that both EGFR and PKA activity were altered in the opposite direction in HOM_Arg943X_ hiPSC-CMs compared to myocardium of obstructive HCM (oHCM) patients and that increased PKA activity further destabilized MTs in HOM_Arg943X_ hiPSC-CMs. Therefore, as a next step, we used the WT hiPSC-CMs to assess whether either kinase is able to modulate the MT code towards a human oHCM-like MT code. As shown in **Figures 8A-D**, in WT hiPSC-CMs, EGFRi mediated an increase of acetylated α-tubulin levels (**Figure S14**). This, again, suggested that EGFR is unlikely to drive the MT densification in HCM myocardium. Thus, we subsequently focused on decreased PKA activity as a driver of increased MT repolymerization in human HCM.

**Figure 8.**
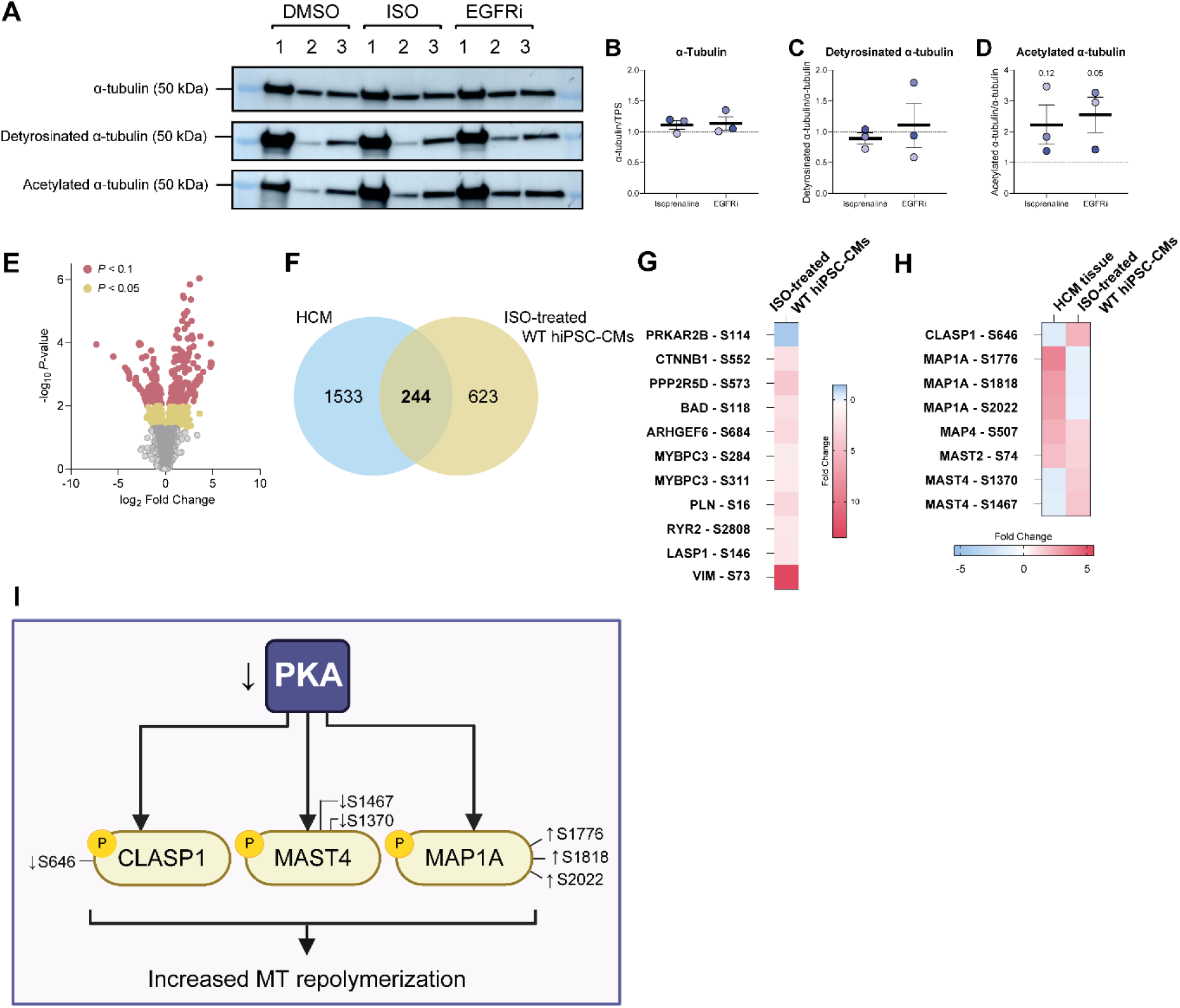
Protein kinase A (PKA) activation in wild-type (WT) hiPSC-CMs using isoprenaline (ISO) leads to differential phosphorylation of microtubule-related and HCM-specific phosphosites. (**A**) Representative blot images of α-tubulin, detyrosinated α-tubulin, and acetylated α-tubulin detection using wild-type (WT) hiPSC-CMs treated with DMSO, ISO and EGFRi. Total α-tubulin (**B**) and detyrosinated α-tubulin levels (**C**) remained unaltered, whereas acetylated α-tubulin levels (**D**) were increased. (**E**) Volcano plot showing significantly altered pSer/pThr sites in ISO-treated WT hiPSC-CMs. (**F**) Venn diagram depicting the phosphoproteomic overlap between the HCM patient and ISO-treated hiPSC-CM datasets. (**G**) All detected phosphosites corresponding to the PKA signaling pathway, which is significantly decreased in HCM patients. (**H**) Heatmap depicting the phosphoproteins shared by the HCM patient and ISO-treated WT hiPSC-CMs and which are involved in modulating MT dynamics. (**I**) Significant phosphosite-specific derived from HCM patient tissue and their overlap with the ISO-treated WT hiPSC-CM-derived phosphotargets. HCM-related fold-changes are in comparison to non-failing myocardium, whereas the fold changes depicted for the hiPSC-CMs are relative to DMSO-treated hiPSC-CMs.

To validate which phosphoproteins downstream of ISO mediated a decreased MT repolymerization, we performed another pSer/pThr screen. To this end, we used 30-days old WT hiPSC-CMs treated with either DMSO or ISO (N=4). We identified 992 pSer/pThr (*P*<0.05, **Figure 8E**) sites that were differentially phosphorylated upon ISO treatment of WT hiPSC-CMs. 244 of these altered (phospho)proteins showed overlap with our previous HCM screen (**Figure 8F**). Importantly, using ISO-treatment, we found hyperphosphorylation and detection of PKA-related phosphosites (**Figure 8G**) which we reported to be hypophosphorylated in HCM (**Figure 3E**). We then checked for overlapping phosphosites that are specific to MTs. As depicted in **Figure 8H**, we showed that upon ISO treatment, *CLASP1* (S646), *MAP1A* (S1776, S1818, and S2022), *MAP4* (S507), *MAST2* (S74), and *MAST4* (S1370 and S1467) overlapped with our HCM myocardial screen. Of these overlapping sites, only *CLASP1*, *MAP1A* and *MAST4* presented with a phosphorylation change directly opposite of HCM tissue. Since HCM patients present with decreased PKA activity, we annotated these (phospho)proteins as potential downstream targets (**Figure 8I**).

In addition to the above, we observed that S22456 on titin, which correlated with both detyrosinated and acetylated α-tubulin levels and was less phosphorylated in HCM patient myocardium, was more phosphorylated in ISO-treated WT hiPSC-CMs. Moreover, we found that ISO decreased the phosphorylation of S895 on LIM domain only protein 7 (LMO7), a site that was hyperphosphorylated in HCM and correlated with diastolic function (e’). Downstream of ISO treatment and linked to clinical parameters, we further found *Abl* (S17, correlated with IVSi), *MAST4* (S1467, correlated with LADi) and *POPDC2* (S333, correlated with LVOTg) to overlap with HCM myocardium. Thus, by focusing on PKA, we were able to further validate a subset of our correlative screen.

## Discussion

We assessed both mRNA transcript and protein levels of the key enzymes mediating MT acetylation and detyrosination in human HCM myocardium. In addition, we analyzed the phosphoproteome and kinome of NF donor and HCM myocardium of different genotypes. We then introduced a novel hiPSC-CM (HOM_Arg943X_) model for HCM and utilized this model to assess the etiological position of MT stability in HCM pathophysiology. Finally, we integrated our findings to dissect the involvement of EGFR and PKA in modulating MTs.

### The HCM phosphoproteome in relation to both cardiac function and MTs

Our data reveals that neither mRNA transcript nor protein levels of key MT modifying enzymes account for the altered MT code in human HCM. Given its essential position in cardiomyocyte contractility through kinase activity, phosphorylation may provide a clue as to whether these key enzymes, then, are subject to HCM-specific PTMs. Our unique phosphoproteomics dataset highlights that HCM myocardium has a distinct phosphoproteomic and kinase activity landscape and adds to previous studies by including Tyr kinase signaling. Although we were unable to detect phosphorylation sites corresponding to the known enzymes mediating MT acetylation and detyrosination, we show that Tyr kinase signaling is increased in the HCM kinome. This is strengthened by enriched signaling of the EGFR, INSR/IGF1R and ERK-MAPK pathways in HCM. Moreover, we found decreased PKA signaling in HCM myocardium. By correlating distinct phosphosites and kinases to comprehensive demographic and clinical datasets, we sought to provide more insight into the kinase signaling pathways in HCM.

To our knowledge, we are the first to determine both mRNA and protein levels of the enzymes upstream of MT acetylation and detyrosination. In failing myocardium, targeting protein levels of TTL or VASH has been an important tool for ameliorating relaxation deficits in cardiomyocytes [15–17, 20, 25]. Based on our analyses in oHCM myocardium, it is important to take into account that changes in MT stability cannot be entirely attributed to levels of these enzymes. Our patient myocardium presents with a large variety in VASH1 levels, for example, which do not correlate with their respective genotypes, whilst TTL levels remain unaltered. In addition, we were unable to detect these enzymes, as well as HDAC6 and αTAT1, in our phosphoproteomics screen. Phosphorylation of these enzymes is not well understood, even though it may regulate their enzymatic activity. For example, TTL phosphorylation is regulated by protein kinase C (PKC) and, with the exception of decreased activity of its ε subunit (PRKCE), we found no changes in the activity of PKC [43]. As of yet, nothing has been reported on the detyrosinating VASH-SVBP complexes. On the other hand, upstream of MT acetylation, phosphorylation of HDAC6 on S22 by GSK3β was previously suggested to enhance its deacetylating capacity. We, however, find increased GSK3β activity alongside increased HDAC6 levels and increased MT acetylation [44]. In addition, phosphorylation of αTAT1 by CDK1, PKA and CK2, is thought to localize it to the cytoplasm, thereby allowing it to acetylate MTs [45]. As we find decreased PKA signaling in HCM, this raises the question whether αTAT1 is also hypophosphorylated and sequestered to the nucleus, which would be in line with its decreased levels. Alongside this, the role of HDAC6, which was also reported to deacetylate sarcomere proteins, and the function of MT acetylation in HCM need to be studied further [22, 46, 47].

We used INKA analysis to score kinase activity in our HCM myocardium and combined this with a PTM-SEA analysis to identify disease-relevant kinases. In doing so, we show that EGFR signaling may be enriched in HCM, as exemplified by our INKA analyses and multiple enriched EGFR-related PTM-SEA pathways (EGFR, EGF and EGFR1), in addition to downregulation of the erlotinib pathway. The involvement of this pathway has already previously been shown in HCM mouse models [13, 48]. Similarly, we find that increased activity of IGF1R, INSR, MAPK1 and MAPK3 align with the enrichment of INSR and ERK1/MAPK3 pathways in HCM. In line with the latter finding and previous reports, we find that kinases (FGFR1 and MAPK14) and pathways involved in cardiac hypertrophy (AKT1 and rapamycin) are enriched in HCM, whereas β-adrenergic receptor signalling is downregulated [49–52]. ERK-MAPK signaling can also be linked to MT changes, as hypoactivity of this network was previously shown to destabilize MTs and raises the question whether its activation in HCM is a driving mechanism [53]. Lastly and to our knowledge, the etiological pathomechanism of several hyper- and hypoactive kinases we report remains unclear or unknown. Using a kinase inhibitor screen, we have previously demonstrated that inhibition of a subset of these kinases improves relaxation of HCM mouse cardiomyocytes [13]. Therefore, these kinases are potential novel targets for relaxation deficits of the heart.

### HOM MYBPC3_Arg943X_ hiPSC-CMs: a novel platform to functionally dissect the MT code in HCM

HCM patient hearts typically present with hypercontractility and diastolic dysfunction. Despite the altered MT code being a key characteristic of patients with oHCM and heart failure, we have a limited understanding of its etiological position due to the limited availability of early disease models. To address this gap, we present a unique hiPSC model carrying both the HCM-associated *MYBPC3*_Arg943X_ variant and a *TUBA1B*-RFP tag. These hiPSC-CMs showed a hypercontractile phenotype alongside abnormal calcium handling, despite a trend towards increased PKA activity at 30 days compared to WT cardiomyocytes. Strikingly, as HOM *MYBPC3*_Arg943X_ reached 60 days after differentiation, they exhibited a prolonged calcium transient that potentially counterbalances the hypercontractility. These changes correspond to the cellular phenotype shown previously in patient-derived iPSC-CMs carrying the *MYBPC3*-associated mutation c.772G>A, which presented with both accelerated cross-bridge dynamics and a prolonged calcium transient [10, 54]. Moreover, the introduction of a frameshift variant into exon 2 of *MYBPC3* was also shown to increase sarcomere shortening of iPSC-CMs with a TTN-GFP tag, but also increased their relaxation time [55]. The normal relaxation time of HOM *MYBPC3*_Arg943X_ iPSC-CMs could reflect an early mutation-specific feature in 2D culture, as patients carrying the Arg943X variant can present with diastolic dysfunction without LV hypertrophy [4]. Alternatively, 30-days old HOM *MYBPC3*_Arg943X_ iPSC-CMs exhibit an increased RT/CT, which may be a proxy for a sarcomere relaxation deficit.

By tracking MT-related changes in a 2D HCM model for the first time, we show that the presence of an HCM genotype mediates more repolymerizing MTs at both 30-and 60-days after start of differentiation. This increased MT repolymerization develops in the absence of consistent changes in MT PTMs compared to WT. In light of this, our findings of key MT enzymes in 2D and in human myectomy tissue show that they are unlikely to act as key regulators in HCM. It is, therefore, probable that an increased MT stability precedes increases of MT detyrosination and acetylation. Such MT PTMs appear to be accumulated instead. In fact, although we show that the inhibition of detyrosination decreases MT repolymerization in 60-days old WT and HOM *MYBPC3*_Arg943X_ hiPSC-CMs, we also show that the increase of MT acetylation in WT hiPSC-CMs has no direct consequence on MT repolymerization. Moreover, although previous studies demonstrate that MT detyrosination and acetylation can impair cardiomyocyte contractility, we found no correlation between the contractile parameters of our HOM *MYBPC3*_Arg943X_ hiPSC-CMs and their MT codes (data not shown).

### The complexity that shapes the MT landscape in cardiomyocytes

Our non-canonical hits from the HCM myocardial screen suggest that the MT and clinical landscape in HCM might not be regulated by one, but multiple kinase signaling pathways. Indeed, we were able to validate the involvement of EGFR and more so, PKA, in the MT landscape of cardiomyocytes. Though EGFRi did not result in a hiPSC-CM phosphoproteome that directly opposed that of HCM myocardium, we do provide evidence that modulating EGFR increases MT acetylation. More strikingly, we show that PKA activation decreases MT repolymerization in HOM *MYBPC3*_Arg943X_ hiPSC-CMs. Given its prominent role as a hypoactive kinase in both HCM and heart failure, this highlights beta-adrenergic receptor signaling as a therapeutic target. We also denote that if therapy is tailored towards MT-modifications, ISO-related downstream (phosphor)proteins such as MAP1A, MAST4 and CLASP1 represent a novel set of treatment targets.

We performed our correlative studies to determine whether clinical parameters associate with distinct kinases and/or phosphosites in our HCM patient population, using the clinical data set of our HCM population. We found that in HCM, EPHA4 is hyperactive. EPHA5-Y833 is also more phosphorylated in HCM, which may be downstream of the hyperactive kinase FGFR1. There is a trend towards increased phosphorylation of Y544 of EPHA2 as well, though its upstream kinase is unknown. Secondly, PKA is upstream of PRKD2 phosphorylation, but none of these sites were detected in our screen. PRKACG, PRKACB, PRKCI, BRAF and MARK1 all correlate to clinical markers, but pathway-specific changes in their phosphosites were not detected. Overall, our screen suggests that EphA and PKA may be more involved in mediating cardiac dysfunction.

### Study limitations

In addition to identifying differences between HCM and NF donors, we sought to compare genotypes, sexes and IVSi (N≥8 per group) within our HCM patient group. We could not find distinct phosphoproteomic differences when categorizing samples according to these covariates. However, previous phosphoproteomic studies using HCM myocardium did find these covariates to alter the HCM phosphoproteome [56–58]. Genotype and sex, for example, have been highlighted as potential players [57, 59, 60]. This discrepancy may arise on a biological level due to study design differences (i.e. ages at the time of myectomy and different head-to-head comparisons). In light of this, it is important to note that we compare HCM myocardium at a progressed disease state to NF myocardium.

Whilst our dataset allows for finding relevant targets at a later disease stage, it also highlights the necessity for phosphoproteome studies in models of early stage HCM, as was achieved before by using hiPSC-derived cardiomyocytes carrying a mutation in the *MYH7* gene [58]. As such, a seemingly decreased EGFR and increased PKA activity in our 30-days old HOM *MYBPC3*_Arg943X_ hiPSC-CMs could be indicative of an early disease stage. Nevertheless, our hiPSC data should be interpreted in light of the immaturity of 2D models, as exemplified by the expression of ssTnI. Further validation of the involvement of PKA, for example, ought to be validated using 3D models such as engineered heart tissue. An important additional consideration is that biological replicates of hiPSC-CMs, generally numbered as separate differentiations, are likely to show variations in MT-related protein despite carrying the same genotype. Therefore, it is also important to study as many replicates as possible, though this finding also underscores that the MT code is more intricately regulated than what was known before.

## Conclusion

In conclusion, our study represents a comprehensive analysis of the pTyr and pSer/pThr residues within HCM myocardium. We provide valuable insight for incorporating the role of phosphorylation in HCM studies, especially those involving clinical and MT data. Importantly, we highlight that EGFR/IGF1R-MAPK signaling and PKA signaling are increased and decreased, respectively, and that PKA is a key regulator of MT repolymerization. Together with protein analyses of the enzymes that mediate MT PTMs and our *MYBPC3*_Arg943X_ hiPSC-CMs studies, we provide a resource for studies focusing on the etiological position of the MT code in HCM and heart failure pathology.

## Acknowledgements

We would like to thank the Ellison Stem Cell Core at the Institute for Stem Cell and Regenerative Medicine (University of Washington) for supporting the generation of the cell lines described in this manuscript.

## Sources of Funding

This work was funded by the Netherlands Cardiovascular Research (CVON) and Dutch CardioVascular Alliance (DCVA) initiatives of the Dutch Heart Foundation (2020B005 DCVA-DOUBLE-DOSE; 2014-40 CVON-DOSIS), a ZonMW VICI grant (91818602) and a Leducq Foundation award 20CVD01. This work was also supported by the NIH HL128368 grant to MR.

## Disclosures

All authors declare that the research was conducted in the absence of any commercial or financial relationships that could be construed as a potential conflict of interest.

## Data availability statement

The data underlying this article will be shared on reasonable request to the corresponding author.

## Notes

### Competing Interest Statement

The authors have declared no competing interest.

